# Mutations in CalDAG-GEFI Lead to Striatal Signaling Deficits and Psychomotor Symptoms in Multiple Species Including Human

**DOI:** 10.1101/709246

**Authors:** Jill R. Crittenden, Magdalena Sauvage, Takashi Kitsukawa, Eric Burguière, Carlos Cepeda, Véronique M. André, Matthias Canault, Morgane Thomsen, Hui Zhang, Cinzia Costa, Giuseppina Martella, Veronica Ghiglieri, Karen A. Pescatore, Ellen M. Unterwald, Walker Jackson, David E. Housman, S. Barak Caine, David Sulzer, Paolo Calabresi, Michael S. Levine, Christine Brefel-Courbon, Anne C. Smith, Marie-Christine Alessi, Jean-Phillipe Azulay, Ann M. Graybiel

## Abstract

Syndromes caused by mutations in Ras-MAP kinase signaling molecules are known as RASopathies and share features such as developmental delay, autistic traits, and cancer. Syndromic features of Rap-MAP kinase signaling defects remain undefined. CalDAG-GEFI is a calcium-responsive Rap-GTPase activator that is enriched in the matrix of the sensorimotor striatum and down-regulated in Huntington’s disease. We show here that CalDAG-GEFI mutations, including striatum-specific deletions and spontaneous mutations in the enzymatic domain, are associated with psychomotor phenotypes in humans, dogs and mice. The identification of these neural mutants was guided by the overt bleeding phenotype in CalDAG-GEFI knockout mice, and then in humans and other species with conserved platelet signaling deficits. Knockout mice exhibit loss of striatal long-term potentiation and deficits in dopamine, acetylcholine and glutamate signaling, along with delayed motor learning and drug-induced perseverative behaviors. Thus, loss of CalDAG-GEFI signaling produces an evolutionarily conserved syndrome characterized by bleeding and psychomotor dysfunction.

## INTRODUCTION

Upon binding to calcium, the guanine nucleotide exchange factor (GEF) activity of CalDAG-GEFI (CDGI) targets Rap-family small G proteins that subsequently drive integrin-adhesion to surface ligands, vesicle release, and mitogen-activated (MAP) kinase activation in hematological cell types (Bergmeier et al., 2007; Crittenden et al., 2004; Ghandour et al., 2007; Kawasaki et al., 1998; Lozano et al., 2016; Niemz et al., 2017). We found that CDGI is strikingly enriched in the striatum (Kawasaki et al., 1998), a key node in the basal ganglia. This observation led us to examine striatal levels of CDGI in Parkinson’s disease and Huntington’s disease, disorders in which striatal abnormalities are foundational etiologic factors for motor and mood symptoms. We found strong down-regulation of CDGI expression in post-mortem tissue from the striatum of Huntington’s disease patients (Crittenden et al., 2010) and in rodent models of both Huntington’s disease and Parkinson’s disease (Crittenden et al., 2009; Crittenden et al., 2010). The direct impacts of CDGI down-regulation in the striatum have, however, remained untested.

To fill this gap, we engineered mice with selective CDGI deletion in the striatum and mice with constitutive CDGI deletion. Alongside a detailed study of the CDGI-deficient mice, we performed the first neurologic and cognitive assessments of humans with deleterious CDGI mutations. We further tested the generalization of the behavioral phenotypes in CDGI mutants by capitalizing on a loss-of-function CDGI mutation known in Basset Hounds (Boudreaux et al., 2007). We report here behavioral abnormalities in the humans, rodents and canines with blocked CDGI signaling, including motor and psychomotor features in each species. In the mouse mutants, we tested for molecular and cellular phenotypes in the striatum that could underlie these neurologic symptoms. We found deficits in striatal plasticity, including loss of striatal long-term potentiation (LTP), and deficits in dopamine and acetylcholine signaling. We conclude that CDGI mutations disrupt cell-surface receptor signaling pathways in both striatal projection neurons and in platelets to produce a syndrome with bleeding and psychomotor symptoms that, with species-specific variations, are evolutionarily conserved from mouse to human.

## RESULTS

### Genetic Engineering in Mice and Spontaneous Mutations in Humans and Hounds Block CDGI Function

We engineered lines of mice with a constitutive or Cre-dependent stop codon near the 5’ end of *CDGI* (Figure S1). In brain tissue from the global knockout mice, we found *CDGI* DNA and mRNA with the expected deletions and loss of CDGI protein expression (Figures 1A-1F and Figure S1). To enable explicit testing of CDGI function in the post-natal striatum, we crossed CDGI^flox/flox^ mice to a mouse line in which striatal Cre activity begins after birth (Lemberger et al., 2007) (Figure 1G).

**Figure 1.**
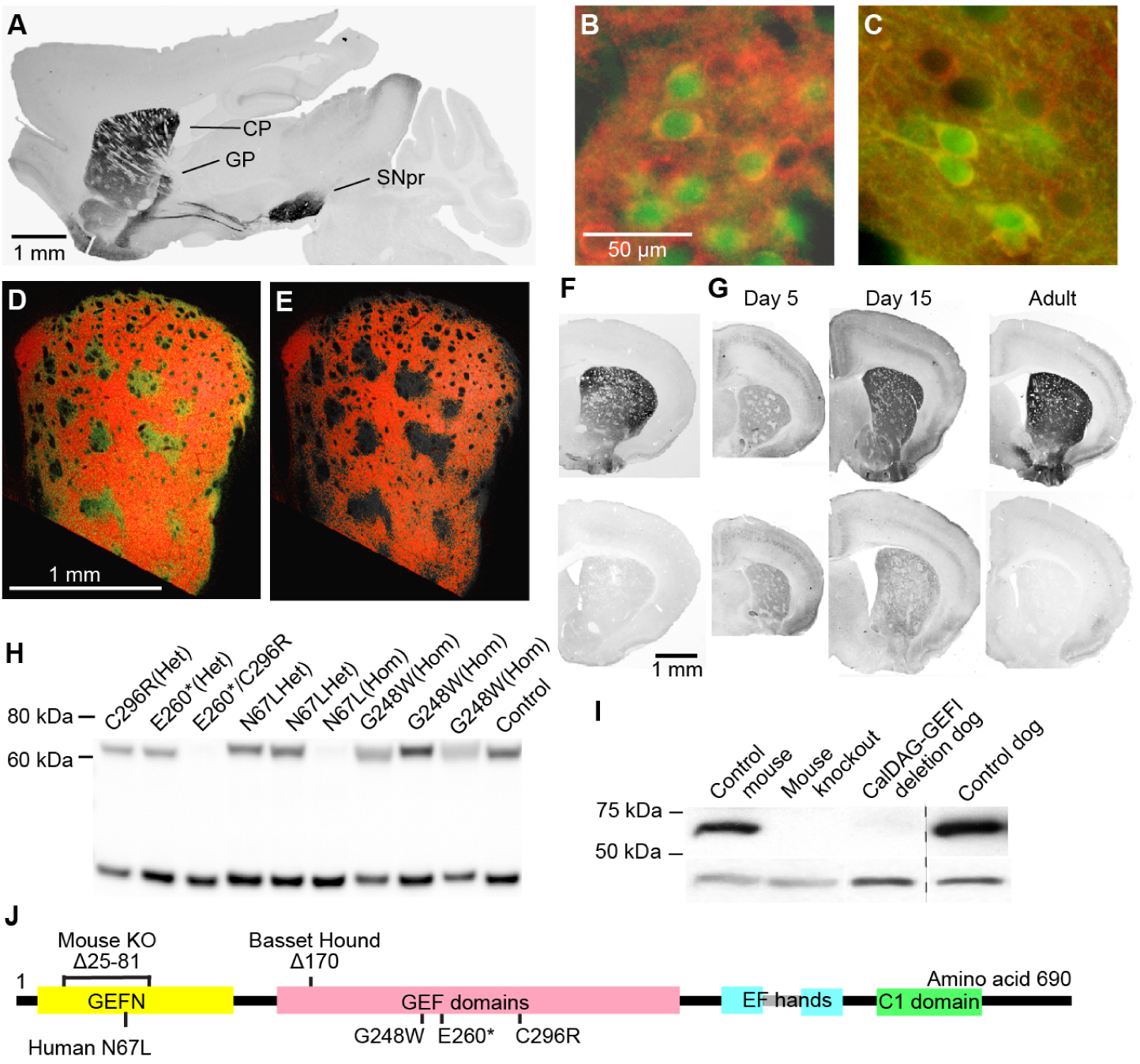
CDGI Is Enriched in Projection Neurons of the Striatal Matrix, and Is Lacking in Mice, Dogs and Humans with Different CDGI Mutations. (A) CDGI immunostaining in sagittal section through mouse brain showing expression in direct and indirect striatal output projections from the caudoputamen (CP) to the external globus pallidus (GP) and substantia nigra pars reticulata (SNpr). (B and C) CDGI immunofluorescence (red) shows co-expression with green fluorescent protein (GFP) in D1 BAC (B) or D2 BAC (C) mice. (D and E) CDGI (red) enriched in matrix, not in striosomes, shown in transverse mouse striatal section co-labeled for the striosomal marker CalDAG-GEFII (CDGII, green channel, shown in (D). (F) CDGI immunoreactivity (black) in control (top) and global knockout (bottom) coronal mouse brain hemisection. (G) CDGI expression is lost postnatally in conditional CDGI^flox/flox^ mice that carry a D1-Cre yeast artificial chromosome (YAC, bottom) but not in CDGI^flox/flox^ control mice (top). (H) Immunoblots showing loss of CDGI in platelet lysates from humans with E260*/C296R and homozygous N67L mutations, but no loss of expression with homozygous G248W mutations. (I) Immunoblots showing loss of CDGI in striatal lysates from a CDGI^ko/ko^ mouse and in platelet lysates from a CDGI homozygous mutant Basset Hound. Lower-band loading-controls show immunoreactivity for GAPDH (H) and beta-actin (I). (J) Diagram of the CDGI protein amino acids showing the mutations tested in mouse, dog and human. See also Figures S1 and S2 and Tables S1-S5.

We tested for loss of CDGI function and expression in humans discovered to have CDGI mutations based on their presentation with a bleeding phenotype similar to that which we earlier reported for CDGI knockout mice (Crittenden et al., 2004; Sevivas et al., 2018; Westbury et al., 2017). We were able to evaluate individuals with four different CDGI mutations, among the eighteen independent mutations so far identified (Table S1). We confirmed that the enzymatic target of CDGI, Rap1, exhibited reduced activation in platelets taken from all of the humans that carried bi-allelic CDGI mutations (Figure S2 and Table S2), as previously reported in CDGI-deficient mice and in a subset of the human patients (Crittenden et al., 2004; Westbury et al., 2017). Thus, all of these mutations represent loss-of-function variants of CDGI. With the same C-terminus antibody used to confirm deletion of CDGI in the mice (Crittenden et al., 2004), we found that two of the three mutation combinations in the human patients resulted in reduced CDGI expression (Figures 1H and 1J): a homozygous frameshift mutation (N67L) and a double heterozygous condition with a premature stop codon in one allele and a missense mutation in the other allele (E260*/C296R) (Westbury et al., 2017) appeared to abolish CDGI protein expression. By contrast, we discovered that the missense mutation G248W within the GEF domain did not block CDGI expression, thereby providing a way to evaluate the consequences of specifically disrupting CDGI enzymatic activity. These results defined a cohort of human subjects with which to examine neurologic functions in relation to discoveries in mice harboring engineered deletions of CDGI.

We further analyzed how CDGI expression is impacted by a 3 base-pair deletion that codes for a conserved amino acid in the catalytic domain of CDGI known in Basset Hounds (Boudreaux et al., 2007). CDGI protein expression was severely down-regulated, but not completely blocked, by the mutation (Figures 1I and 1J) as assessed for the first time with a knockout-validated antibody. Together, these results lay the foundation for cross-species analysis of how loss-of-function CDGI mutations affect behavior and striatal function.

### CDGI Mutations Are Associated with Abnormal Striatum-Based Behaviors in Mice

We began by examining global CDGI knockout mice (Figure S3 and Tables S3-S6) and found that they were fertile and that they had normal gross brain morphology and expression of striatal protein markers, normal transcriptome-wide striatal mRNA expression levels, normal levels of total striatal amino acids and biogenic amines and normal dopamine receptor binding in the striatum. There were no differences between CDGI knockouts and sibling controls in adult weight, daily chow consumption, rotarod motor coordination, open field behavior, olfactory acuity, responses on a SHIRPA (Rogers et al., 1997) neurological exam, bouts of defined gross motor activities measured in a 24-hour home-cage scan of the mice, marble-burying behavior or social-recognition memory. Thus, basic behaviors appeared normal in mice with a constitutive knockout deletion in CDGI.

In tests of striatum-based learning, however, the CDGI mutant mice were impaired. We employed a pair of maze tasks designed to assess differential striatal and hippocampal learning strategies (mainly egocentric and allocentric, respectively) (Packard and McGaugh, 1996). Control mice rapidly acquired an egocentric T-maze task, but CDGI knockouts exhibited significantly delayed learning, (Figure 2A). The CDGI knockout mice showed no deficits in the allocentric maze task, or in assays of amygdaloid complex function (Figure S3).

**Figure 2.**
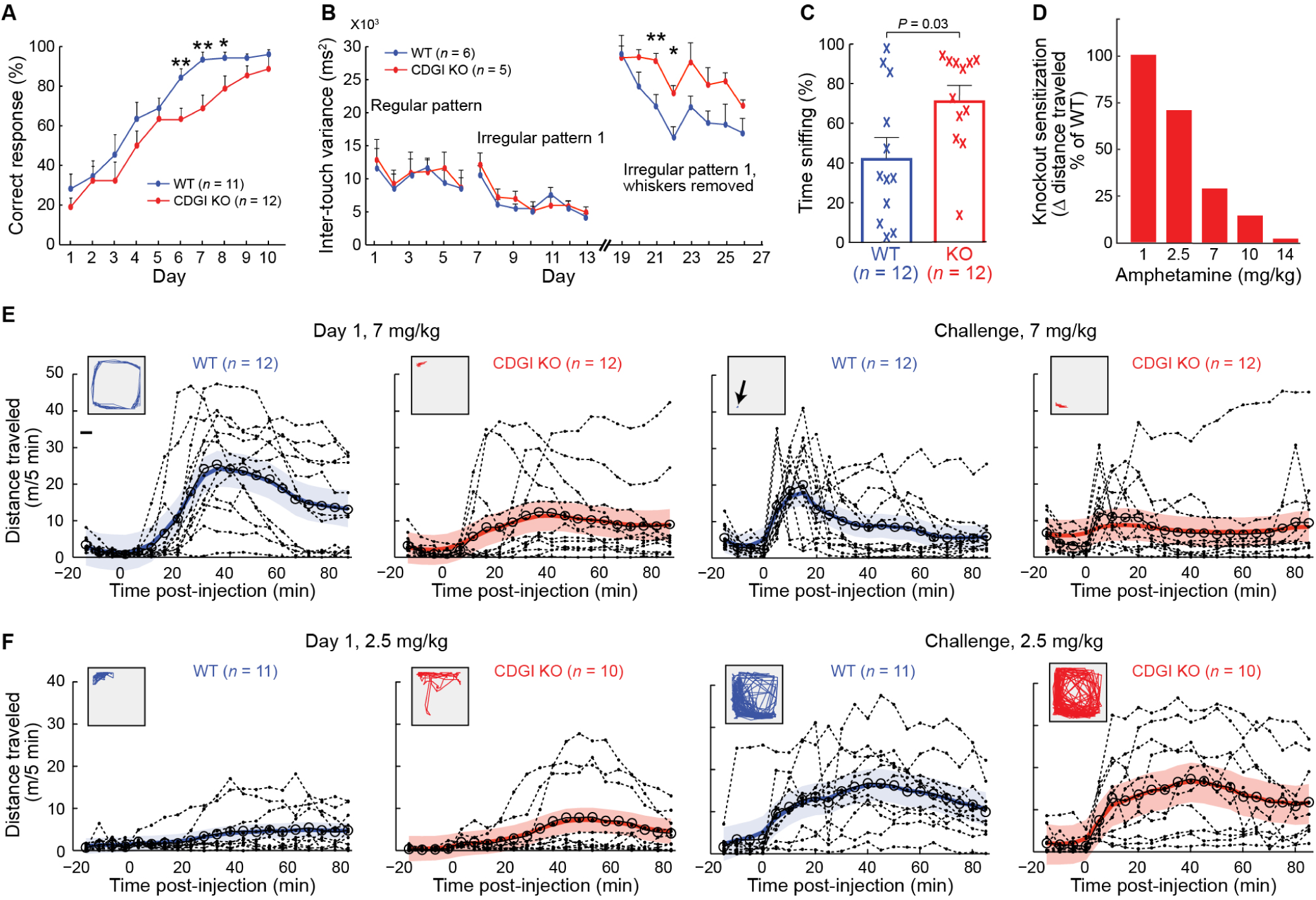
Global CDGI Knockout Mice Exhibit Deficits in Assays for Striatal Learning. (A) In an egocentric T-maze task, knockout (KO) mice learned more slowly than wildtypes (WTs; *P* < 0.05 by ANOVA; *P* = 0.009 on day 6, *P* = 0.006 on day 7 and *P* = 0.04 on day 8 by unpaired, two-tailed Student’s t-test). Error bars show standard errors of the mean. (B) KO mice showed delayed learning in a motor sequence task. Mice were trained in a running wheel with unevenly spaced foot-rungs, and learning was measured as a reduction in the variance of paw placement on a designated rung (Nakamura et al., 2017). Intact KOs and controls learned at equivalent rates; after whisker-cutting, KOs showed delayed re-acquisition. \*\**P* = 0.01 and \**P* = 0.02 by unpaired, two-tailed Student’s t-test between genotypes. (C) KOs spent more time than WTs in sniffing stereotypies after acute amphetamine (7 mg/kg). *P* value was calculated by two-tailed Mann-Whitney U test. (D-F) With increasing amphetamine doses, KOs engaged in less locomotion than WTs, consistent with increased stereotypy. Locomotor sensitization to high-dose amphetamine was occluded in KOs (D). Sensitization was calculated as the change in distance travelled on day 1 vs. challenge day at 50-55 min post-injection (stereotypy-response period). *n =* 10WT/10KO, 11WT/10KO, 12WT/12KO, 4WT/4KO and 8WT/8KO, respectively, for 1, 2.5, 7, 10 and 14 mg/kg of amphetamine. With high-dose amphetamine, WTs showed reduced locomotion on challenge day relative to day 1, consistent with stereotypy sensitization (E). KOs were stereotypic on day 1, so locomotor sensitization was occluded. KOs showed normal sensitization to low-dose amphetamine as shown by an increase in distance travelled after repeated treatments (challenge day, right panels in F) relative to the first day of treatment (left panels in F). Dotted lines represent raw data from each mouse; large open circles are population means; colored lines are random effects estimates of the median with 90% confidence intervals. Insets show sample open-field tracker plots (50-55 min post-injection). See also Figures S3-S5 and Table S6.

We next tested the knockout mice on a sensitive peg-wheel running assay of striatal learning (Kitsukawa et al., 2011; Nakamura et al., 2017) in which the arrangement of the wheel’s left and right footstep pegs could be changed to test the ability of the mice to learn different running patterns (Figures 2B and S3). Learning was measured as reduced variance in the timing of paw-placement on the pegs. CDGI knockout mice performed complex peg-running tasks as well as their sibling controls. When we clipped their whiskers, however, challenging their sensory feed-back, the CDGI knockouts learned the task more slowly, as measured by a higher variance in timing of paw-placement relative to that of whisker-clipped controls. The unmasking of a motor sequence learning phenotype in the CDGI knockout mice upon sensory deprivation is consistent with a deficit in the processing of sensorimotor inputs that are known to favor the matrix compartment (Figures 1D and 1E) in the dorsolateral striatum (Kincaid and Wilson, 1996; Moussa et al., 2011; Packard and McGaugh, 1996). It is in this striatal compartment in which CDGI is highly concentrated in the mouse striatum (Kawasaki et al., 1998).

**Figure 3.**
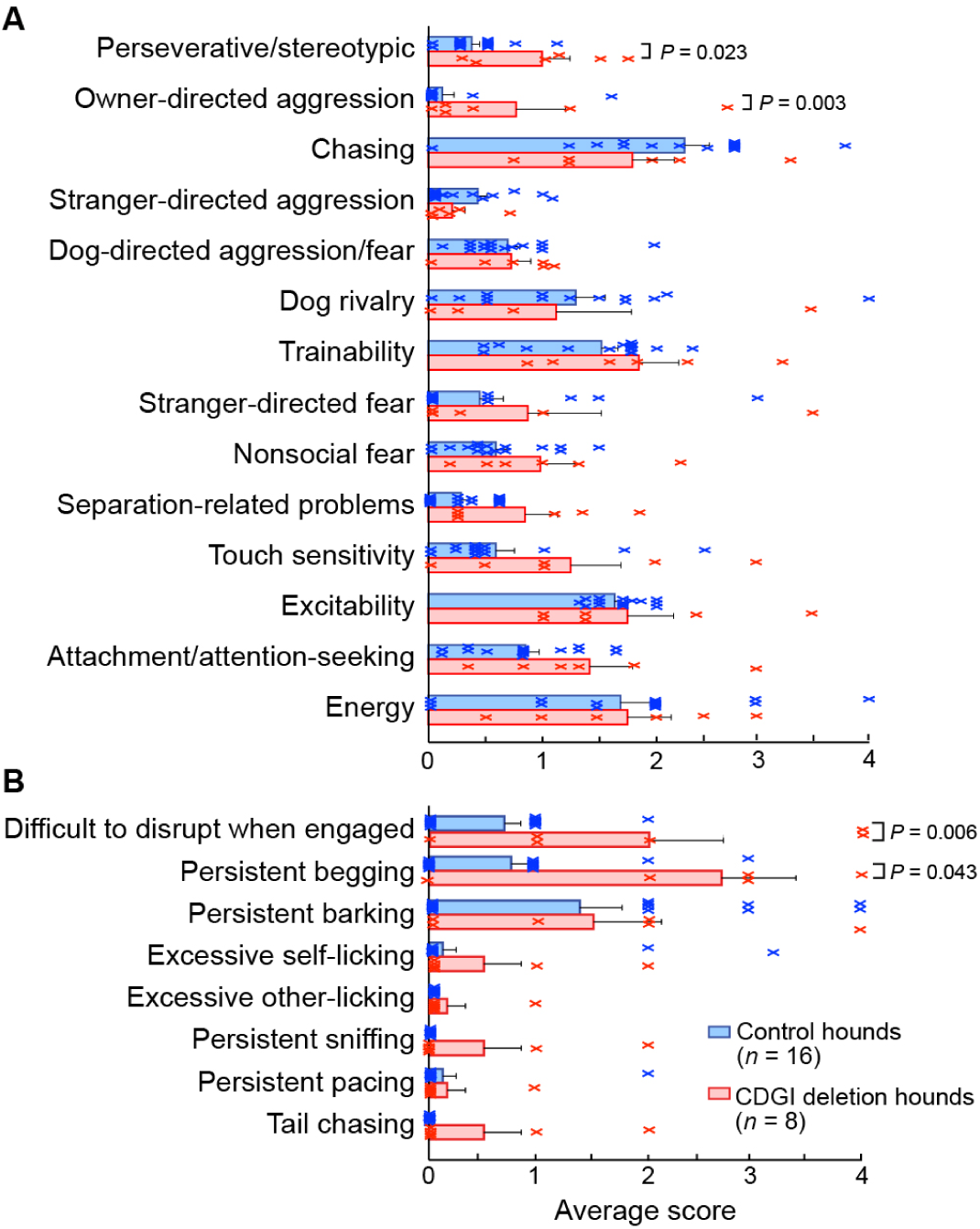
Compiled Behavioral Questionnaire Responses from Owners of Basset Hounds Lacking CDGI Expression and from Control Basset Hounds. (A) Dog-owners’ responses on the 105-question survey were divided into the 14 pre-defined behavioral categories that are listed to left; the scores (ranging from 0 to a maximum of 4) were then averaged. Hounds with CDGI mutations scored higher in two categories: perseverative/stereotypic behaviors and owner-directed aggression. P values were calculated by two-tailed Mann-Whitney test. Error bars show standard error of the mean. (B) Within the perseverative/stereotypic behaviors category, scores were significantly higher for hounds with CDGI deletions, relative to controls, in queries of persistent begging and difficulty to disrupt when engaged.

We then took a clue from the fact that animals exhibit confined, stereotypic behavioral responses to high or repeated doses of psychomotor stimulants such as amphetamine, but not to low doses, and that these behaviors reflect signaling in the dorsal striatum (Canales and Graybiel, 2000; Crittenden and Graybiel, 2017; Kuczenski and Segal, 1997). CDGI knockouts given high doses of d-amphetamine showed greater levels of localized, focused stereotypies and less distance traveled than the wildtype controls (Figures 2C, 2E, S4 and S5) on the very first day of drug treatment. CDGI knockouts failed to exhibit the drug sensitization shown by wildtypes in response to a final drug-challenge dose given after repeated drug treatments and a drug washout period (Figures 2D, 2E, S4 and S5). Similar results were obtained in CDGI^flox/flox^ mice with Cre-mediated postnatal deletion (Figures S4 and S5). These results indicated that CDGI acts to oppose drug-induced stereotypy and overly focused behavior.

The abnormal amphetamine response phenotype in the knockouts did not appear to result from differential drug metabolism or from differential sensitivity to the neurotoxic effects of repeated amphetamine injection, as judged by levels of serum amphetamine or total striatal dopamine and its metabolite, homovanillic acid (HVA), measured after amphetamine injection on the challenge day (Figure S4). Measurements of distance traveled after injection of D1- and D2-type dopamine receptor agonists showed no genotype differences (Figure S4), suggesting that direct downstream signaling from D1 and D2 dopamine receptors is intact in CDGI knockout mice. These findings could indicate that the abnormal response of the mice to amphetamine was produced by a circuit defect upstream or parallel to the dopamine receptor system.

### Humans with CDGI Mutations Exhibit Signs Found in Humans with Basal-Ganglia Related Circuit Disorders

We evaluated behavior in four men and five women aged from 19 to 61 years who carry CDGI mutations (Figures 1H and 1J and Table 1). Of the nine individuals examined, four were heterozygous and five were homozygous or double trans-heterozygous for a mutation in CDGI (Canault et al., 2014). All of these mutations are within highly conserved domains of the protein that are important for its exchange factor activity (Figure 1J), domains that we targeted for deletion in the mouse.

**Table 1.** Individuals with Deleterious Mutations in CDGI Show Symptoms Consistent with Basal Ganglia Dysfunction.

| **Patients family/ subject** | **Genetics** | **Sex** | **Age** | **Occupation** | **Medical background** | **Treatments** | **Behavior** | **Movement disorders** | **Addictions** | **Cognition dysexecutive functions** |
| --- | --- | --- | --- | --- | --- | --- | --- | --- | --- | --- |
| **1.1** | Homozygous G248W | F | 61 | Accountant | Hepatitis<br>Tuberculosis<br>High blood pressure | Perindopril | Insomnia<br>Anxiety | Chorea<br>Both upper limbs | 0 | Yes (short term and visuo spatial memory impairment, planification)<br>Self-reported memory problems |
| **1.2** | Homozygous G248W | M | 59 | Disability | HIV<br>Neuropathy | Anti-HIV | Insomnia | Ataxia | Tobacco,<br>Cannabis | Yes (short term and visuo spatial memory impairment)<br>Self-reported memory problems |
| **1.3** | Homozygous G248W | M | 55 | Engineer | 0 | 0 | Insomnia<br>Anxiety | Self-reported fasciculations | Tobacco,<br>Alcohol,<br>Cannabis | Yes (deficit of working memory, cognitive flexibility, attention and slowness) |
| **2.1** | Heterozygous C296R | M | 47 | Engineer | 0 | 0 | Insomnia<br>Anxiety | Chorea<br>Upper right limb | Unknown | Yes (mild dysexecutive syndrome: lexical evocation, perseverations, attention deficit, visuoperceptive dysfunction) |
| **2.2** | Heterozygous E260* | F | 46 | Engineer | 0 | 0 | 0 | None | Unknown | No |
| **2.3** | Double trans-heterozygous C296R E260* | F | 19 | Student | 0 | 0 | 0 | Self-reported tremor when tired | Unknown | Yes (mild dysexecutive syndrome: attention deficit, cognitive flexibility)<br>Absence of memory impairment |
| **3.1** | Heterozygous N67Lfs*2 | M | 50 | Unemployed | Diabetes<br>Coronaropathy | 0 | Insomnia | None | Tobacco,<br>Alcohol | Refused cognitive exam<br>Illiterate |
| **3.2** | Heterozygous N67Lfs*2 | F | 54 | Farmer | Breast Cancer | 0 | Insomnia | Dystonia | Tobacco | Mild deficit in working memory |
| **3.3** | Homozygous N67Lfs*2 | F | 23 | 0 | 0 | 0 | Tiredness | Chorea both lower limbs | Tobacco,<br>Alcohol | Yes (deficit in working memory, programing, lexical fluency)<br>Illiterate |

Of the 9 people examined, 6/9 complained of insomnia, 3/9 complained of anxiety, 7/9 exhibited a movement disorder, 5/6 self-reported use of addictive substances and 6/7 were identified to have dysexecutive symptoms.

The individuals with CDGI mutations exhibited a triad of psychomotor abnormalities associated with basal ganglia dysfunction. Of the five individuals with mutations on both chromosomes who were given neurological exams, three were diagnosed with mild chorea and one with ataxia, movement abnormalities typical of basal ganglia dysfunction. The heterozygotes either received a diagnosis of chorea (*n* = 1) or dystonia (*n* = 1), or were scored as normal on motor exam (*n* = 2). Notably, six out of the seven individuals who completed the cognitive tests were scored as having mild executive dysfunction mainly affecting working memory, cognitive flexibility, attention and slowness, despite the fact that two were engineers without apparent professional difficulties. Five of the six individuals queried about addictive substance use confirmed that they use tobacco, alcohol or cannabis. Finally, seven of the nine individuals complained of insomnia, tiredness and/or anxiety. No abnormal repetitive behaviors were reported, paralleling our findings in untreated CDGI knockout mice.

The abnormalities in these individuals were mild, but nonetheless they indicate that humans with mutations affecting the enzymatic domain of CDGI have a syndromic disorder consisting of not only an increased tendency for bleeding due to an evolutionarily conserved failure in integrin-mediated platelet adhesion, but also motor, cognitive and behavioral abnormalities.

### CDGI Mutations Are Associated with Abnormal Behaviors across Species

With veterinarians, we identified a group of Basset Hounds that were either homozygous for a CDGI mutation (*n* = 8) that disrupts CDGI protein expression (Figures 1I and 1J) or confirmed not to have the mutation (*n* = 16) (personal communication, Dr. M. Boudreaux). The owners (*n* = 10) were given under single-blind conditions a C-BARQ behavioral questionnaire (Hsu and Serpell, 2003) modified by the insertions of eight questions related to the overly focused, repetitive behavioral phenotype of drug-treated CDGI knockout mice. Of the 14 behavioral domains assessed, CDGI mutants were rated as significantly different from non-mutant hounds in only two of the categories queried: ‘perseverative behaviors’ including ‘difficulty to disrupt when engaged’ and ‘persistent begging’ and ‘owner-directed aggression’ (Figures 3A and 3B).

This pre-clinical evidence added to the indications of abnormal psychomotor behavior in CDGI deficient humans and engineered mice, and indicated that mutations blocking CDGI activity produce a syndrome of species-modulated behavioral abnormalities typically associated with basal ganglia dysfunction, and an accompanying bleeding disorder, across species.

### Spontaneous Activity and AMPA and NMDA Currents Are Abnormal in Striatal Projection Neurons of CDGI Mutant Mice

We explored possible underlying signaling defects in the striatum of CDGI mutants by examining the activity of the dominant class of striatal neurons, the spiny projection neurons (SPNs, also called medium spiny neurons), which normally are enriched for CDGI (Figures 1A-1C). We recorded their spontaneous activity and responses related to glutamatergic, dopaminergic and cholinergic transmission, three key modulators of SPN activity.

Whole-cell patch clamp recordings of SPNs in slices from CDGI knockout mice and sibling controls indicated equivalent membrane capacitance, input resistance and time constants (Table S7). By contrast, recordings of synaptic currents demonstrated a significantly greater frequency of spontaneous excitatory post-synaptic currents (EPSCs) in striatal slices from CDGI knockout mice than from controls under baseline conditions and with bath-applied bicuculline, a GABA receptor blocker, and with 4-aminopyridine, a potassium channel blocker that promotes neurotransmitter release (Figure 4A). These results suggested an increase in presynaptic glutamate release in the striatum of CDGI knockouts. In addition, there were significant changes in the kinetics of spontaneous synaptic events. Half-width and decay times were significantly reduced in SPNs from CDGI knockouts compared to controls (Figure 4B), and average amplitude differences tended to be higher but this trend failed to reach significance (−9.97 ± 0.45 pA in controls and −11.25 ± 1.23 pA in knockouts).

**Figure 4.**
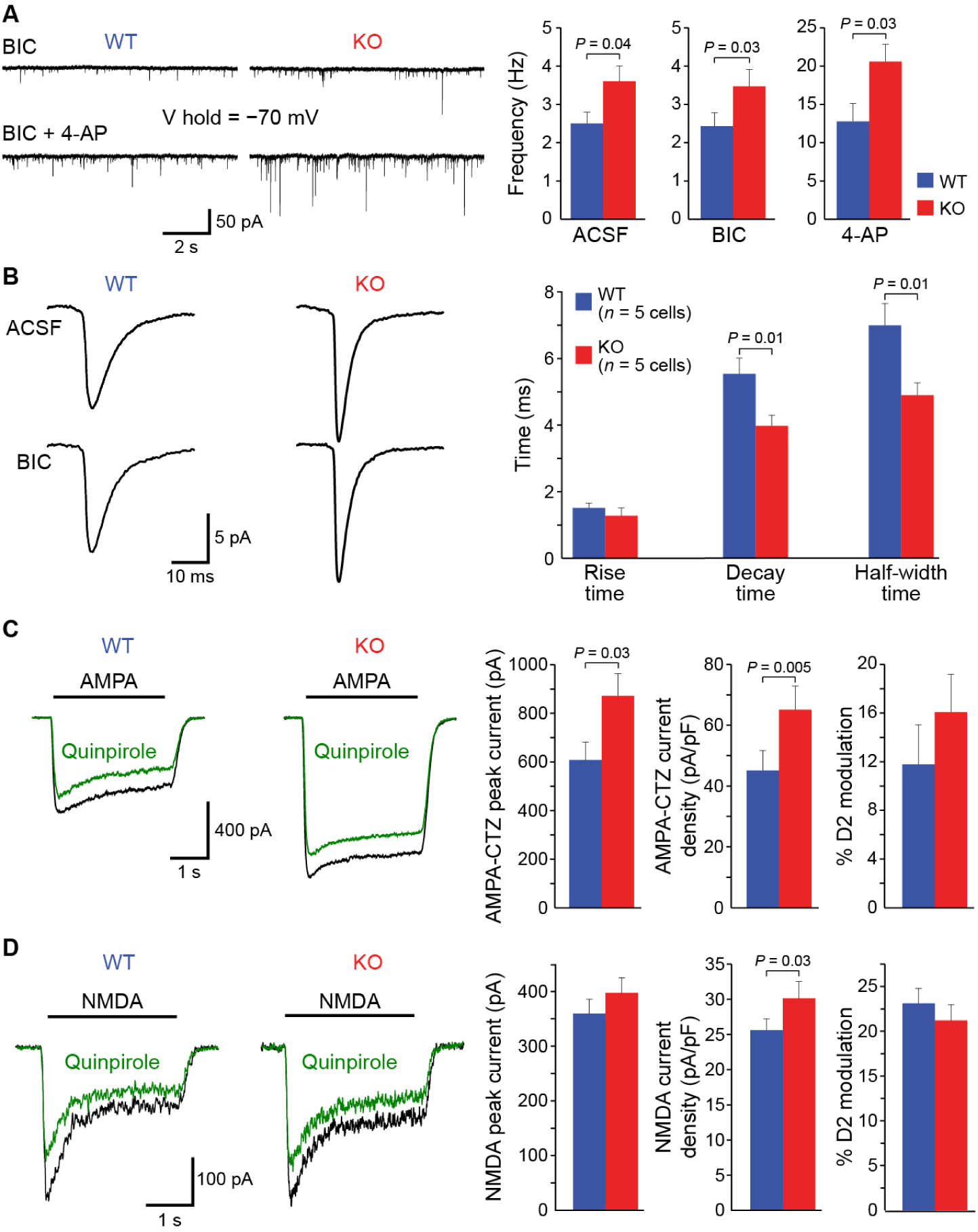
Global CDGI Knockouts Exhibit Increased Spontaneous and Evoked AMPA and NMDA Receptor Currents in SPNs. (A) Spontaneous EPSCs recorded in SPNs of striatal slices after addition of bicuculline (BIC, 10 µM) alone or with 4-aminopyridine (4-AP, 100 µM) in artificial cerebral spinal fluid (ACSF). Average frequency data are shown at right. *n* (cells) = 16WT/23KO for ACSF, 16WT/23KO for BIC, and 12WT/12KO for 4-AP. Error bars show standard error of the mean. (B) Average spontaneous EPSCs (5-50 pA) in ACSF and after addition of BIC show faster kinetics in SPNs from KOs. Decay time and half-amplitude duration were significantly shorter in cells from KOs, relative to WTs. (C) Current responses of dissociated striatal neurons to AMPA application (100 µM) in the presence of cyclothiazide (CTZ, 10 µM) to prevent receptor desensitization. Bar graphs show significant increases of both peak AMPA current and AMPA current density in KOs (*n* = 24), relative to WTs (*n* = 12). Application of the D2 dopamine receptor agonist, quinpirole (10 µM), reduced AMPA current to a similar degree in cells from WTs (*n* = 9) and KOs (*n* = 19). (D) Current responses to NMDA (100 µM) application. Average peak NMDA currents were not significantly different between groups, whereas the current density was increased in cells from KOs following normalization by cell capacitance (*n* = 15WT, 29KO). Application of the D2 dopamine receptor agonist, quinpirole (10 µM), reduced NMDA current to a similar degree in cells from WTs (*n* = 6) and KOs (*n* = 8). Statistical comparisons made by two-tailed Student’s t-test. See also Tables S7-S8.

We next recorded from acutely dissociated striatal neurons to isolate postsynaptic responses to AMPA and NMDA glutamate receptor activation. Basic membrane properties were normal in dissociated SPNs from CDGI knockouts (Table S8). However, average peak currents and current densities of the AMPA-mediated responses were significantly greater in the knockouts than in wildtypes (Figure 4C). The peak NMDA-mediated current was equivalent between genotypes, but current density was significantly greater in knockouts, following normalization for cell capacitance (Figure 4D). These findings suggest that, at least *in vitro*, SPNs in CDGI mutants exhibited abnormally high activity both spontaneously and in response to glutamatergic stimulation.

### Dopamine Signaling Is Abnormal in the Striatum of CDGI Mutant Mice

Dissociated SPNs from the CDGI knockout mice exhibited normal reductions of both AMPA- and NMDA-mediated currents in response to the D2 dopamine receptor agonist, quinpirole (Figures 4C and 4D). D1-type dopamine receptors were not examined for technical reasons, but these results at least suggest normal post-synaptic D2-type dopamine receptor modulation of excitatory SPN currents in CDGI knockouts.

However, with *in vivo* microdialysis in awake, resting CDGI mutants and controls, extracellular dopamine levels in the striatum of the knockouts were half those of the wildtypes (Figure 5A). There was also a trend for reduced striatal levels of the dopamine catabolites dihydroxyphenylacetic acid (DOPAC) and HVA. We tested whether these diminished levels of striatal dopamine reflected enhanced dopamine catabolism via catechol-O-methyl transferase (COMT). We found equivalent COMT activity in striatal tissue from knockouts and controls, after repeated saline or repeated amphetamine (14 mg/kg/day) treatments (Figure S4). Thus, the CDGI knockouts had reduced levels of extracellular dopamine despite normal total striatal dopamine and HVA content and apparently normal dopamine metabolism.

**Figure 5.**
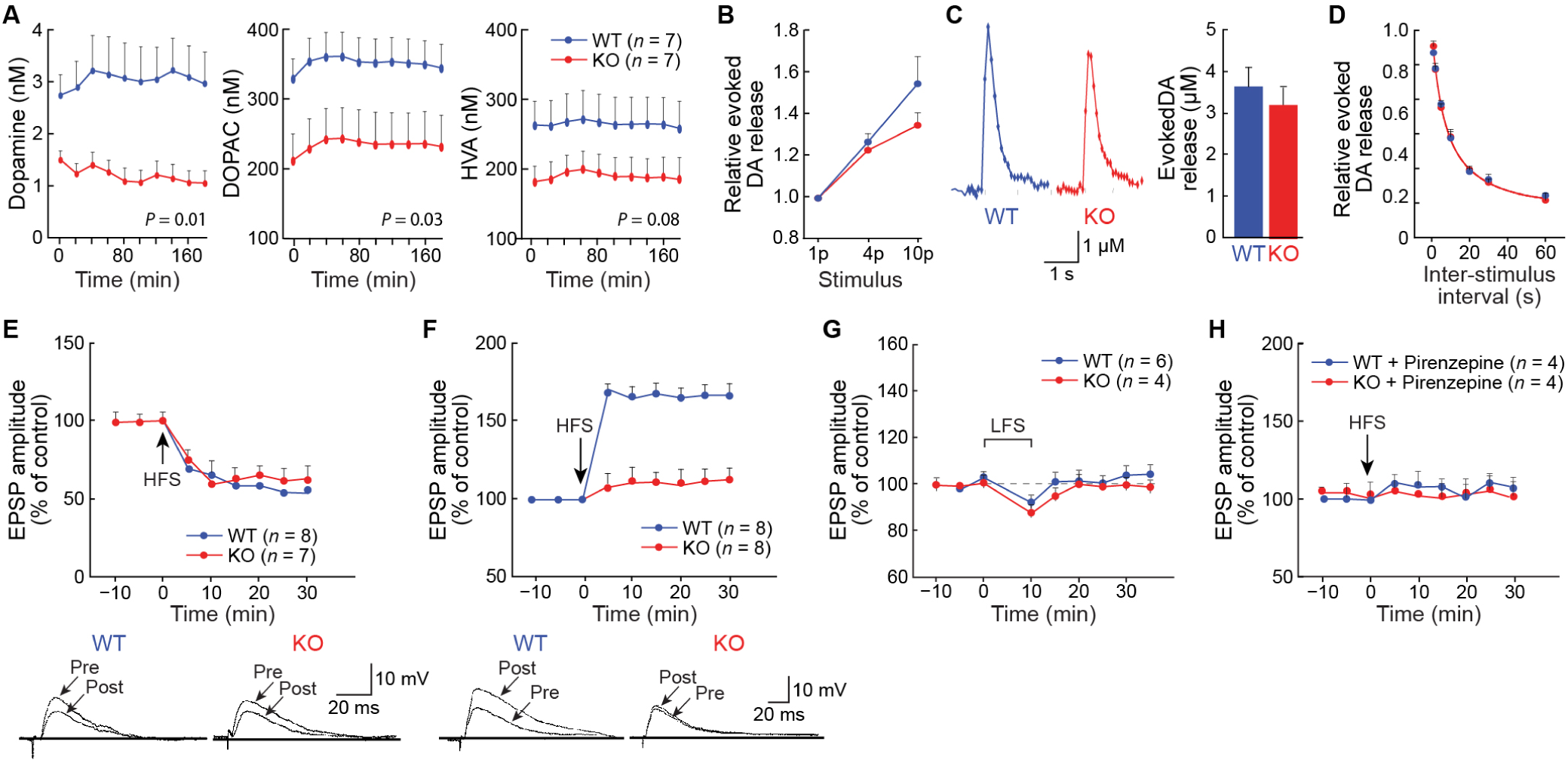
CDGI KO Mice Have Diminished Extracellular Dopamine and Loss of Striatal LTP. (A) *In vivo* microdialysis in the dorsal striatum showed reduced dopamine and DOPAC and a trend for reduced HVA in KOs, relative to WTs. *P* values were calculated by Student’s unpaired, two-tailed t-tests between average values. Error bars show standard error of the mean. (B) Dopamine release in response to a single pulse (1p) or train stimuli (4 or 10 paired stimuli at 100 Hz) was equivalent in KOs and controls. *n* = 14 slices for 1p, *n* = 8 slices for 4p and for 10p with slices taken from 5 mice per genotype per experiment. (C) Examples (left) and averages (right) of dopamine response to 1p. (D) Paired-pulse stimulation showed equivalent depression of evoked dopamine release between genotypes (*P* > 0.05, Student’s two-tailed t-test, *n* = 7 slices from each of 5 mice per genotype). (E) With magnesium, high-frequency stimulation (HFS) induced striatal LTD in slices from both genotypes (*P* < 0.001, one-way ANOVA followed by Bonferroni post-hoc test). (F) With no added magnesium, HFS of corticostriatal synapses induced LTP in WTs (*P* < 0.001), but not KOs (*P* > 0.05). Resting membrane potentials were −84mV (E) and −85 mV (F) for both genotypes. Shown below are examples of neuronal EPSPs recorded immediately before (pre) and 20 min after (post) HFS. (G) WTs and KOs exhibited equivalent responses to low-frequency stimulation (LFS) for depotentiation (*P* > 0.05). (H) The M1 receptor antagonist pirenzepine (100 nM, 5 min preincubation) blocked LTP in WTs and had no effect in KOs (*P* > 0.05 by two-tailed Student’s t-test for paired data).

Finally, we used fast-scan cyclic voltammetry in striatal slice preparations (Figures 5B-5D) to test for dopamine release evoked in response to single stimuli and trains of stimuli designed to emulate phasic firing (4 and 10 pulses of 100 Hz). The release with single stimuli was equivalent in the knockouts and controls. The release with trains of stimuli was nearly equivalent between genotypes, but with 10 pulses, there was 30% lower release in knockouts than in wildtypes. Paired pulse depression was normal in CDGI knockouts at inter-pulse intervals ranging from 1 to 60 sec. Together, these findings indicate that evoked dopamine release was normal or slightly reduced in the CDGI knockouts, suggesting that the deficiency in baseline extracellular levels of striatal dopamine could arise from a circuit-level failure to excite the dopamine-containing neurons or their axonal release sites rather than from factors intrinsic to the state of the terminals.

### Measurements of LTP and LTD Demonstrate Loss of LTP in the Striatum of CDGI Mouse Mutants

Signaling through dopamine and NMDA receptors is key for corticostriatal plasticity (Calabresi et al., 1992; Partridge et al., 2000; Shen et al., 2008) that is suggested to be an electrophysiological correlate of behavioral plasticity (Centonze et al., 2006; Kreitzer and Malenka, 2008). We tested for the presence of these mechanisms in CDGI knockout mice and their littermate controls. In this preparation, we found similar resting membrane potential, input resistance and responses to slow depolarizing ramp voltage commands and current injections (Figure S6). Stimulation of cortical afferents induced equivalent long-term depression (LTD) in the dorsolateral striatum in brain slices taken from CDGI knockouts and sibling controls, as measured by intracellular recordings of excitatory post-synaptic potentials (EPSPs) in SPNs (Figure 5E). SPNs in slices from wildtype mice showed normal LTP when input fibers were stimulated in the absence of magnesium, which blocks NMDA receptors (Calabresi et al., 1992). By contrast, equivalent stimulation in striatal slices from knockout mice failed to induce LTP (Figure 5F).

We considered the possibility that the striatal LTP deficit in the knockouts resulted from saturation of striatal LTP in these mice (Centonze et al., 2006). Such slices are refractory to LTP induction, but they exhibit depotentiation with a protocol that can reverse LTP. We subjected striatal slices from the knockouts and wildtypes to low-frequency stimulation of cortical fibers, which reverses striatal LTP (Centonze et al., 2006), but found equivalent responses in striatal slices from knockouts and wildtypes (Figure 5G). Thus, SPNs from CDGI knockouts were not in a state of saturated LTP.

Striatal activity and LTP are strongly modulated by the M1 muscarinic acetylcholine receptor (Calabresi et al., 1999; Perez-Burgos et al., 2010; Shen et al., 2007), which is expressed in SPNs and other striatal cell types. In heterologous cell culture assays, CDGI can signal downstream of the M1 muscarinic receptor (Guo et al., 2001). In our experiments, LTP induction in wildtypes was blocked by the M1 inhibitor, pirenzepine (Figure 5H) as expected (Calabresi et al., 1999), and application of pirenzepine to slices from CDGI knockout mice did not change their lack of response to LTP induction protocols. Thus, the blockade of M1 receptors in the wildtypes mimicked the loss of LTP phenotype from CDGI genetic deletion. These findings are consistent with CDGI signaling downstream of the M1 muscarinic receptor in SPNs to mediate corticostriatal LTP.

### CDGI Knockout Mice Exhibit Deficits in Behavioral Modulation by the Muscarinic M1 Receptor

Given the repetitive behaviors in the hounds and drug-treated CDGI mutant mice, the established links between repetitive behaviors and M1 receptor signaling in the dorsal striatum (Aliane et al., 2011; McCool et al., 2008), and the *in vitro* link of M1 cholinergic receptors to CDGI signaling, we examined CDGI knockout mice in a cocaine self-administration assay in which self-administration can be blocked by the specific M1 receptor allosteric agonist, VU0357017-5 (Thomsen et al., 2010). Firstly, there were no differences between genotypes in nose-poking behavior when the nose-pokes were reinforced by administration of liquid food reward (not shown) or by escalating doses of intravenous cocaine (Figure 6A). As expected, when the wildtype sibling mice were treated systemically with VU0357017-5, they exhibited a reduction in cocaine self-administration (Figures 6A and 6B). By contrast, the CDGI knockout mice continued to self-administer cocaine after VU0357017-5 treatment, indicating that they were insensitive to M1 modulation of self-administration. When saline replaced cocaine in the self-administration procedure, the CDGI knockout mice exhibited the normal extinction of nose-poking behavior (Figures 6A and 6B). These findings suggest that CDGI normally mediates signal transduction from the M1 muscarinic receptor in SPNs, and that this signaling could mitigate excessive motoric responses to dopamine receptor signaling, including signaling augmented by psychomotor stimulants.

**Figure 6.**
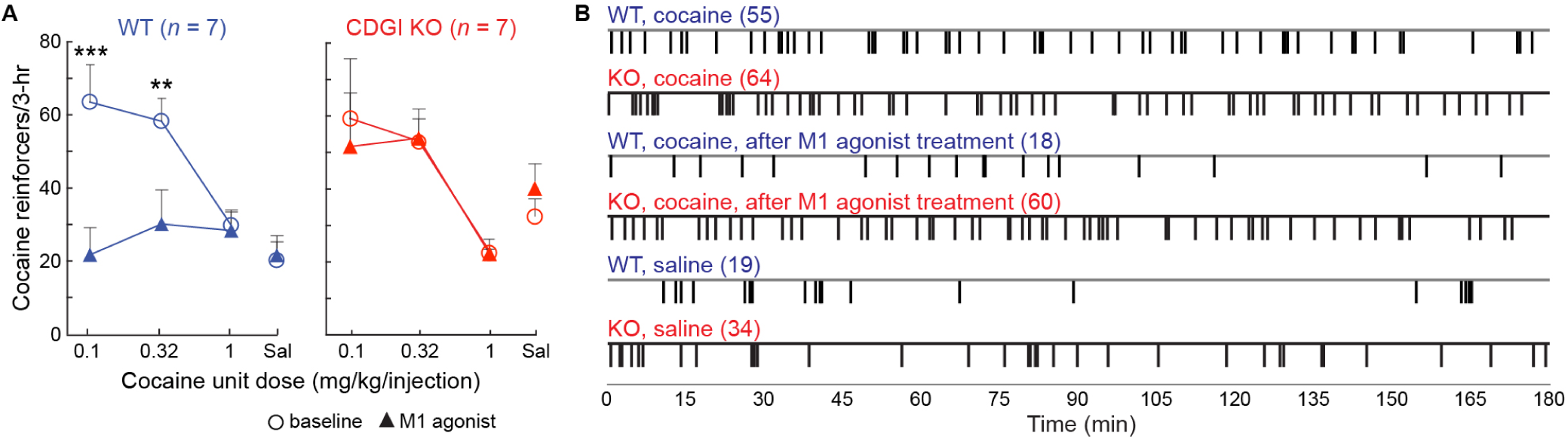
Global CDGI Knockout Mice Fail to Show Normal Suppression of Drug Administration in Response to M1 Acetylcholine Receptor Stimulation. (A) WT and KO mice exhibited dose-dependent self-administration of cocaine (dose, F(3,18)=6.67, *P* = 0.003 and F(3,18) = 6.76, *P* = 0.003, respectively). The M1AchR allosteric agonist VU0357017 suppressed self-administration of cocaine in WTs (left) but not in KOs (right; treatment, F(1,6)=9.52, *P* = 0.02; treatment-cocaine interaction, F(3,18) = 9.29, *P* = 0.0006). \*\**P* < 0.01, \*\*\**P* < 0.001 vs. cocaine alone with Bonferroni post-test. (B) Examples of sessions from WT and KO mice showing each nose-poke (tick-marks) for self-administration of cocaine (0.32 mg/kg/infusion), cocaine after VU0357017 treatment and saline after extinction of cocaine delivery. The numbers in parenthesis correspond to the total number of reinforcers earned in the session.

## DISCUSSION

The striatum is placed in a crucial position within basal ganglia circuitry, as it receives massive inputs from the neocortex and thalamus, and targets its outputs mainly to the pallidonigral nuclei that modulate movement control. Much of this input-output circuitry involves the large matrix compartment of the striatum, which accordingly is viewed as the principle sensorimotor component of the striatum, as opposed to the limbic connections of the striosomes. The predominant expression of CDGI in SPNs in the matrix compartment of the striatum led us to expect that CDGI knockout mice might mainly exhibit sensory-motor deficits if indeed the CDGI protein were important to SPN signaling. CDGI deletions did have behavioral consequences, but contrary to expectation, we found little evidence for primary sensory or motor deficits in the knockout mice, with either global or striatum-restricted CDGI deletion. Instead, we found deficits in behavioral learning and plasticity under challenging conditions, across cognitive as well as motor domains, and a loss of LTP, a cellular marker of synaptic plasticity. These findings suggest that the matrix compartment as a whole might perform computations engaged by the motor and action demands of complex environments. Our surveys of humans and canines that lack CDGI expression or function showed that they, too, have psychomotor phenotypes, albeit in species-modulated forms. Three of the humans in our study did not show loss of CDGI expression, but did show loss of CDGI’s ability to activate Rap1, combined with motor and cognitive features that overlapped with those found in the other families. Our findings thus suggest that deficient CDGI enzymatic activity is critical to the expression of a syndrome characterized by platelet dysfunction and bleeding as well as striatal dysfunction with psychomotor abnormalities.

Our evidence for the presence of CDGI in the striatum is confirmed by mRNA and protein analyses in the mouse and human (Crittenden et al., 2010), and traits compatible with striatal dysfunction (Boraud et al., 2018; Graybiel, 2008) were present upon neurologic examination of the human subjects: hyperkinetic movement disorders including mild chorea, tremor and dystonia, behavioral manifestations including anxiety, insomnia and drug use, and a mild dysexecutive syndrome. Platelet aggregation responses were clearly better in heterozygous than in biallelic mutant individuals, but the neurologic symptoms were more variable among the genotypes. Based on the phenotypes in CDGI mutant mice and canines, we tested for obsessive-compulsive behaviors in these individuals but did not find these. Nevertheless, of the nine individuals with heterozygous or biallelic mutations in CDGI, all but one (heterozygote for the E260* mutation) showed motor and/or cognitive problems.

We cannot consider that any of these individuals have an overt neurologic disease, but the clinical manifestations that we observed are present in major basal ganglia disorders such as Parkinson’s disease and Huntington’s disease. Our findings here suggest that the down-regulation of CDGI, which occurs in Huntington’s disease (Crittenden et al., 2010) and in models of L-DOPA-induced dyskinesia (Crittenden et al., 2009), might contribute to the chorea and dystonia that occur in these disorders. Moreover, the impacts of CDGI loss might be more evident under disease or stress conditions, as behavioral abnormalities in the mice were only uncovered upon psychomotor drug treatment and under challenging learning paradigms. These findings suggest that the functions of CDGI across multiple organ systems are conserved across species and produce conjoint hematopoietic and striatal defects that underlie bleeding and behavioral phenotypes.

We were able to test striatal signaling in mouse, but not in human or hound. However, the experiments in our mouse model were highly informative. Glutamate signaling was abnormal as tested in dissociated SPNs and in striatal slices from the mutants. The frequency of spontaneous EPSCs was increased, suggesting presynaptic alterations. The kinetics of these synaptic events were abnormal for both AMPA and NMDA receptor signaling. These findings hint at the possibility that thalamic or neocortical inputs to striatal neurons, or both, could be abnormal in CDGI mutants.

Finally, we found a total absence of corticostriatal LTP in CDGI mutant slice preparations, without detectable changes in corticostriatal LTD. This striking finding indicates that there is abnormal plasticity in corticostriatal or thalamostriatal circuits, or other intrinsic basal ganglia circuits impacting striatal function (Mallet et al., 2012) possibly related to the delayed sensorimotor learning that we found in the CDGI mutant mice. NMDA receptor function in the dorsolateral (sensorimotor) striatum, which is particularly enriched for CDGI expression, is essential for normal learning on the wheel-running assay that we used (Nakamura et al., 2017). Thus, the NMDA signaling abnormalities in the striatum of CDGI mutants might directly underlie their slowed learning in this task. The fact that CDGI knockouts were normal in numerous other memory and motor tests further suggests that the defects occurring in the absence of CDGI signaling are functionally specific.

Baseline extracellular dopamine content in the striatum of CDGI knockout mice was half that of the wildtype levels measured in microdialysis experiments, and behavioral responses to indirect dopamine agonists were abnormal, as manifested by severely confined and repetitive behaviors. Nevertheless, evoked dopamine release as tested *in vitro* with fast-scan cyclic voltammetry was nearly normal in CDGI mutants. A working hypothesis to account for these findings is that circuit-based dysfunction occurs in the CDGI mutants. This circuit abnormality could possibly occur through abnormal anterograde or retrograde SPN signaling that, in turn, controls dopamine terminals or dopamine-containing neurons themselves to produce diminished baseline dopamine levels and hyper-responsivity to high doses of psychomotor stimulants.

The sensorimotor and stereotypic symptoms that appear with mutation of CDGI are similar to those following disturbances of basal ganglia circuits including nigro-striato-nigral loop-circuits. This pattern is consonant with the fact that CDGI is particularly enriched in the output neurons of the sensorimotor striatum. CDGI immunoreactivity fills the cell bodies and fine processes of D1- and D2-dopamine receptor-positive SPNs within the striatal matrix, which receives preferential input from sensorimotor and prefrontal cortices (Kincaid and Wilson, 1996; Ragsdale and Graybiel, 1990), and in the D1- and D2-positive SPN output pathways (Kawasaki et al., 1998). By contrast, there is lower expression of CDGI in striosomes and in the ventral striatum (Kawasaki et al., 1998), both of which are considered to be related to limbic functions. CDGI knockout mice exposed to psychomotor stimulants showed nearly normal locomotor responses and initial self-administration, but they exhibited exacerbated stereotypy and failed to stop self-administering cocaine in response to M1 muscarinic receptor agonists. Thus, CDGI signaling defines a circuit that distinguishes locomotor responses from stereotypic responses, overly focused behaviors and severe habitual responses. These psychomotor abnormalities resemble aspects of symptoms in autism-spectrum and attention-deficit disorders (American Psychiatric Association, 2013).

Mice with perturbations in cholinergic signaling show exacerbated stereotypic responses to amphetamine and cocaine, as did the CDGI mutants, and they maintain relatively normal locomotor responses (Crittenden et al., 2014; Janickova et al., 2017). Such drug-induced stereotypic behavior is known to be regulated by dopamine and acetylcholine signaling in the dorsal striatum (Aliane et al., 2011; Capper-Loup et al., 2002; Kuczenski and Segal, 1997, 2001). The pattern of cholinergic neuropil in the striatum is similar to that of CDGI for being abundant in the dorsal striatal matrix compartment (Crittenden et al., 2014; Crittenden et al., 2017; Dautan, 2014; Graybiel and Ragsdale, 1978). Moreover, in a cell culture system, the M1 muscarinic receptor was shown to induce CDGI signaling to Rap1 and the MAP kinase cascade (Guo et al., 2001). Thus, CDGI expression in SPNs is placed to transduce signals from cholinergic interneurons to basal ganglia outputs that affect specific behaviors.

Our findings leave important issues to be resolved. CDGI functions in relation to the immune system (Crittenden et al., 2004; Ghandour et al., 2007), but we did not further explore this system here. The sample sizes for our study of the humans and hounds were constrained, and our data on the hounds were based on surveys. Although consistent with basal ganglia dysfunction, the features exhibited by the different species were individual to the species, insofar as we were able to observe them, and testing for corticostriatal LTP and other signaling features of mutants was only feasible in the engineered CDGI knockout mice. The profound loss of corticostriatal LTP in the mice is of special interest for further work on striatum-based learning and neuroplasticity across species. Given the possibilities for compensation of the effects of CDGI deletion, it is notable that CDGI is necessary not only for normal platelet function, but also for normal psychomotor expression and motor function as tested here. Other CDGI mutations, both in humans and other animals, have been reported. The effects of these mutations, not studied here, could expose the full range of CDGI functions based on the observed CDGI expression in the developing and mature brain and cells of the hematopoietic system. Our findings do however, introduce a novel syndromic disorder produced by mutations in a Rap-MAP kinase activating factor.

## ACKNOWLEDGMENTS

We thank Dr. James Serpell and Dr. Joseph Garner for advice on how to conduct the behavioral survey with dog owners, Dr. Mary Boudreaux for Basset Hound genotype and owner-contact information, Dr. James Catalfamo for the gift of blood platelets from control and mutant hounds, Dr. Charlie Whittaker for bioinformatics analyses, Dr. Yasuo Kubota for assistance in manuscript preparation, and Tao Liu, Michael Yim, Patricia Harlan, Hilary Bowden and Kyle Fischer for technical assistance. This work was funded by the National Institute of Child Health and Development (R37-HD028341, A.M.G.), the James and Pat Poitras Research Fund (A.M.G.), The Saks Kavanaugh Foundation (A.M.G.), The Simons Foundation (A.M.G, J.R.C. and D.S.), The Stanley Center for Psychiatric Research at the Broad Institute, via a grant to Edward Scolnick from the Stanley Medical Research Institute (A.M.G. and J.R.C.), the National Institute of Mental Health (R01-MH071847, A.E.S.; F32-MH065815, J.R.C.), the National Institute of Health (R01-AG050548, A.E.S.), the European Community FP7 – Thematic priority HEALTH contract number 222918 (REPLACES) (P.C.), the Ministry of Health Grants (B.P. and P.C.), the JPB Foundation (D.S.) and the National Institute on Drug Abuse (R00-DA027825, M.T.; R0107418, D.S.)

## AUTHOR CONTRIBUTIONS

J.R.C. and A.M.G. conceived of and supervised the project with the assistance of D.E.H. J.R.C. and A.M.G. wrote the paper with the assistance of M.S., T.K., E.B., C.Cepeda., M.C., M.T., H.Z., E.M.U., W.J., D.S., P.C., A.C.S. and J.-P. A. J.R.C. and A.M.G. analyzed the data, with input from all authors. J.R.C., M.S., T.K. and E.B. performed motor behavior and learning and memory assays. J.R.C. and M.S. performed amphetamine-response assays. W.J. performed 24-hr behavior scans. C. Cepeda and V.M.A. performed assays of glutamate receptor currents in slice physiology experiments under the supervision of M.S.L. M.C. performed human platelet assays under the supervision of M.-C. A. M.T. performed cocaine self-administration assays with the assistance and supervision of S.B.C. H.Z. performed fast-scan slice-voltammetry experiments under the supervision of D.S. C. Costa and G.M. performed long-term potentiation in slice physiology experiments under the supervision of P.C. V.G. performed depotentiation in slice physiology experiments under the supervision of P.C. K.A.P. performed the dopamine receptor ligand radiography under the supervision of E.M.U. C. B.-C. and J.-P. A. performed neurological and behavioral assays with the human subjects. A.C.S. evaluated statistical significance of the amphetamine-response behavioral data from mice.

## STAR METHODS

### METHOD DETAILS

#### Human Neurological Exams

All patients in this study were invited to come for consultation in the neurology department by phone. The patients agreed and came to the consultation outside of any hospitalization and on their own initiative. All patients signed a consent form accepting that the results of their neurological examination be used for research purposes. Subjects were given a cognitive evaluation (16-item free and cued recall, RL/RI-16, Rey-Osterrieth figure recall, WAIS-III, trail making test, Stroop test and VOSP battery) and were examined by clinicians (J-P. Azulay and C. Brefel-Courbon) specialized in movement disorders.

#### Human Platelet Activation Assays

All the participants of this study signed a consent form for the examination of genetic characteristics and the use of their samples for research purposes (AC-2017-2986 and AC-2018-3105, authorization provided by the French Ministry of Education and Research and the local Direction for Research and Innovation). This research is part of the French reference center on hereditary platelet diseases coordinated by Prof. M. C. Alessi.

Washed platelets were suspended in Tyrode’s buffer (138 mM NaCl, 2.7 mM KCl, 12 mM NaHCO3, 0.4 mM NaH2PO4, 1 mM MgCl2, 2 mM CaCl2, 5 mM Hepes, 3.5 mg/ml HSA, and 5.5 mM glucose, pH 7.3) supplemented with 0.02 U/ml apyrase. Studies were performed within 6 h after blood collection by placing platelet rich plasma in an aggregometer cuvette at 37°C with stirring. ADP (5 and 10 µM; Sigma-Aldrich) and TRAP-14 (5 and 10 µm; Polypeptide group) were added and light transmission was recorded on an APACT 4004 optical aggregation system (Labor Bio-Medical Technologies GmbH).

Rap1 activation was determined using a commercially available kit (Millipore) as previously described (Canault et al., 2014). Briefly, washed platelets (3 × 108 platelets/ml) were stimulated for 1 min in non-stirring conditions with ADP (10 µM). Reactions were stopped with ice-cold 2x Rap1 lysis buffer complemented with protease inhibitor cocktail (Roche), and phosphatase inhibitors (NaF, 10 mM and Na3VO4, 1 mM; Sigma-Aldrich). The cell lysates were incubated for 45 min with RalGDS-RBD beads to pull-down Rap1-GTP. Washed pellets were solubilized in sample buffer prior western blotting. Individual proteins were detected with a rabbit polyclonal antibody against Rap1 (Millipore) and a secondary goat anti-rabbit HRP-coupled antibody (Bio-Rad Laboratories). Proteins were detected by chemiluminescence. Total Rap1 levels were detected from whole platelet lysates.

#### Mouse Maintenance

All experiments were approved by, and performed in strict accordance with, the Massachusetts Institute of Technology (MIT) Committee on Animal Care, which is accredited by AAALAC. Mice were group-housed and maintained under a standard light/dark cycle with free access to food and water except during food-reinforced learning and memory experiments, in which cases mice were single-housed. For the cocaine self-administration assay (Figure 6) and the total biogenic amine and amino acid data (Tables S4 and S5), CDGI knockouts and sibling controls were in a congenic C57BL/6J genetic background. For the amphetamine-response experiments with conditional, post-natal CDGI knockouts and sibling controls, mice were in a congenic 129S4 genetic background. For all other experiments, mice were in an isogenic 129S4 genetic background. Male and female mice were used for the cocaine self-administration assay. For the remainder of the experiments, which were conducted prior to the National Institute of Health policy for the inclusion of both sexes in experimentation, male mice were used.

#### Generation of CalDAG-GEFI Knockout Mice

The CDGI targeting construct was based on a 6.2 kb SacI restriction fragment from the 129Sv/cJ7 mouse chromosome 19 BAC clone 7D23 (Guru et al., 1999) that was subcloned into Bluescript II KS+ plasmid (Stratagene Inc.). One loxP site was ligated to a HindIII site in intron 4 and a loxP-flanked fusion gene for hygromycin resistance and thymidine kinase was ligated to an AflII site in intron 2. The targeting construct was electroporated into J1 mouse embryonic stem (ES) cells (gift of Prof. Rudolf Jaenisch) that were subsequently grown in hygromycin to select for integration. Resistant ES cell clones were screened for homologous integration by polymerase chain reaction (PCR) and Southern blotting. Clonal populations were transiently transfected with the Cre recombinase vector Pog231 and gancylcovir was applied to select for loss of thymidine kinase. PCR was used to identify deletion clones, two of which were injected into blastocysts by the MIT Department of Comparative Medicine facility. Resulting chimeric mice were crossed to C57BL/6 to test for germline transmission by coat color and transmitting males were crossed to 129S4 mice to establish the mutation in a background isogenic with the J1 ES cells. Phenotypic analyses were always performed on sibling progeny from pure 129S4 heterozygous mutant intercross matings. Constitutive CDGI knockout mice were genotyped by PCR with the following three oligonucleotide primers: 5’–aacagttcccaggctagagatagagagttcctcc–3’, 5’– accagactctaggccagaacctacc–3’, and 5’–agtgtgctgtggtgaaatgcagccattcc–3’. Wildtype mice yielded a 208 base pair product with the first two primers while the knockout yielded a 286 base pair product with the second two primers. Conditional CDGI floxed mice were genotyped by PCR with the two primers 5’– TCTCAGCTAGTCCATTTCCCAACTAGCGAGTTGC–3’ and 5’–AACAGTTCCCAGGCTAGAGATAGAGAGTTCCTCC–3’ to yield a 650 bp product from the floxed allele and a 594 bp product from the wildtype allele.

#### Reverse Transcriptase Polymerase Chain Reaction

Mouse brain RNA was prepared using the RNAeasy kit (Qiagen). RT-PCR was performed using the ThermoScript RT-PCR kit (Invitrogen) with two primers flanking the sites where loxP was inserted: 5’–TAATACGACTCACTATAGGGAGGCTGAGCTGGTTCAAGTG–3’ and 5’– ATTTAGGTGACACTATAGAACTGCCGCTTCCACTTGTAGG–3’.

#### Western Blotting

For mouse striatal and cortical tissue, mice were deeply anesthetized with pentobarbital (150 mg/kg by intraperitoneal injection), and tissue was dissected on a cold plate prior to freezing in liquid nitrogen and storage at −80^°^C. For dog platelets, blood was taken from two control dogs and one Basset Hound confirmed to be a homozygous mutant for the 3-basepair deletion. Washed platelets were prepared by Dr. James Catalfamo in Ithaca, NY, stored at −80^°^C and shipped to MIT for Western blotting. Frozen samples from mice or dogs were homogenized in ice-cold modified RIPA buffer (50 mM Tris pH 8.0, 150 mM NaCl, 1% Triton X-100, 0.1% sodium dodecyl sulfate, 1% NaDeoxycholate) with Complete protease inhibitor cocktail (Roche) and centrifuged at 16,000 × g for 10 min to pellet insoluble material. Protein concentration of supernatants was determined by bicinchoninic acid assays (Thermo Fisher Scientific). Proteins were resolved by SDS-PAGE and transferred to PVDF membrane (Millipore) by electroblotting. Blots were incubated overnight, at 4°C, with antibodies diluted in TBST [10 mM Tris-HCl (ph 8.0), 150 mM NaCl, 0.05% Tween 20)] and 5% bovine serum albumin. Blots were subsequently washed in TBST and incubated with horseradish peroxidase-coupled secondary antibodies (Vector Laboratories) prior to immunodetection with Western Lightening (PerkinElmer Inc.) according to the manufacturer’s instructions. Blots were subsequently incubated with anti-β-tubulin (Cell Signaling Technology) to control for protein loading. Antibodies were obtained from the following companies: total GluR1 receptor (Calbiochem), total ERK1/2, phospho-ERK1/2, total JNK, phospho-JNK, total P38, phospho P38, total DARPP-32 and phospho-DARPP-32 (Cell Signaling Biotechnologies); GluR2 receptor, phosphor-Ser831 GluR1 receptor, mGluR1 receptor, mGluR5 receptor, M1-type muscarinic receptor, D1-type and D2-type dopamine receptor (Chemicon); phosphor-Ser845 GluR1 receptor (Novus Biologicals); Rap1, Rap2, CalDAG-GEFII/RasGRP2 (Santa Cruz Biotechnologies); phosphor-CREB (Rockland Immunochemicals); PSD95 (Upstate Biotechnologies).

#### Immunolabeling

Mouse brain sections were prepared for immunolabeling as previously described (Crittenden, 2017). The sections were incubated with rabbit polyclonal CDGI antiserum (Crittenden et al., 2004) for approximately 12 h at 4°C and processed either for immunohistochemistry with ABC amplification (Vector Labs) and DAB detection with nickel enhancement or, in conjunction with mouse anti-CalDAG-GEFII antiserum (SC-8430, Santa Cruz Biotechnology), for immunofluorescence with secondary antibodies coupled to ALEXA 564 and ALEXA488 (Invitrogen Corp.). Fluorescent labeling of D1 and D2 dopamine receptor-expressing neurons was detected with an EGFP filter set in sections from Drd1a-EGFP and Drd2-EGFP BAC transgenic mice (Gong et al., 2003). Sections were mounted and coverslipped with Eukitt (Electron Microscopy Sciences), following immunohistochemistry, or with Vectashield media (Vector Laboratories), following immunofluorescence. The sections were viewed with Olympus BX61 and SZX7 microscopes fitted with an Olympus DP70 camera.

#### Exon microarray

Striatal tissue was collected in parallel in three different experiments from age-matched male CDGI knockout mice and sibling controls. After dissection, samples were frozen on dry ice and then homogenized in Trizol (Sigma-Aldrich) and total RNA was prepared according to manufacturer’s instructions and previously described methods (Cantuti-Castelvetri et al., 2005). Equivalent amounts of RNA from each sample were pooled according to genotype (*n* = 4 of each genotype pooled for two experiments and *n* = 3 of each genotype pooled for one experiment) and given to the MIT BioMicro center to prepare cDNA for hybridization to the Affymetrix GeneChip® Mouse Exon 1.0 ST. The MIT Bioinformatics core facility used Affymetrix Expression Console software to summarize and normalize data from the chips.

#### Measurements of Total and Extracellular Striatal Dopamine, DOPAC and HVA

For microdialysis, guide cannulae were implanted in the striatum using stereotaxic surgery. After at least one week for surgery recovery, the microdialysis probe (CMA/7 probe 1mm, CMA Microdialysis, Sweden) was lowered into the guide cannula and microdialysates were collected on ice in perchloric acid (0.5 M) at 20-min intervals at a rate of 1.5 μl/min. Samples collected for the first hour were discarded and subsequently collected samples were immediately frozen and kept at –80°C until HPLC analysis.

For measurements of dopamine, DOPAC and HVA to assess COMT activity, mice were decapitated 50 min after saline or amphetamine injection and the striata were dissected on ice and kept at –80°C until shipment to the laboratory of Prof. Tim Maher (Massachusetts College of Pharmacy and Health Sciences) for high performance liquid chromatography (HPLC).

For HPLC analysis, microdialysates were injected unmodified and striatal tissues were homogenized in 0.2 M perchloric acid, 0.2 mM disodium EDTA and 0.2 mM ascorbic acid. Samples were assayed for DOPAC, dopamine and HVA by HPLC with the potential set at +300 mV with respect to a palladium-hydrogen reference electrode.

#### Measurements of Striatal Amino Acids and Biogenic Amines

For measurements of total amino acids and biogenic amines, male mice between 4 and 5 months of age (*n* = 11WT/7KO) were sacrificed by cervical dislocation, and striatal tissue was dissected and frozen on dry ice for shipment to the Vanderbilt Neurochemistry core for HPLC measurements.

#### Dopamine Receptor Autoradiography

D1- and D2-type dopamine receptor autoradiography was performed as described in Unterwald et al. (Unterwald et al., 1994; Unterwald et al., 2001) D3 receptor autoradiography was carried out according to the method described by Guitart-Masip et al. (Guitart-Masip et al., 2006). Mouse brains were stored at –80°C prior to dissection. Mouse brains were mounted on cryostat chucks using embedding matrix, cut on a Reichert and Jung 2800 Frigocut N into 16 µm coronal sections based on the Paxinos and Franklin mouse brain atlas (Paxinos, 2001), thaw-mounted onto Fisher superfrost glass slides, air-dried on ice and stored desiccated at –30^◦^C until assayed. Slide-mounted sections were preincubated in buffer containing 50 mM Tris HCl, 120 mM NaCl, 5 mM KCl, 2 mM CaCl2 and 1 mM MgCl2, pH 7.4, at room temperature for 30 min. Following preincubation, sections were incubated for 45 min at room temperature in the same Tris-salt buffer in the presence of 1 µM mianserin and 5nM ^3^H-SCH23390, without and with 10 µM fluphenazine to measure, respectively, total and nonspecific binding for the D1-type dopamine receptor. For D2-type receptor autoradiography, slides were incubated in the preincubation buffer with 0.001% ascorbic acid, 1 µM mianserin and 5 nM ^3^H-raclopride without and with 10 µM (+)butaclamol to measure total and nonspecific binding, respectively. For D3 receptor binding, slides were incubated in the preincubation buffer in the presence of 0.001% ascorbic acid, 5 nM ^3^H-PD128907 without and with 1 µM (+)butaclamol to measure total and nonspecific binding respectively. After incubation, slides were washed twice in the Tris-salt buffer on ice for 5 min/wash followed by rinse in ice-cold distilled water. Sections were dried under a cold air stream and allowed to sit overnight at room temperature. Slides for dopamine receptor autoradiography and tritium standards (Amersham) were exposed to tritium-sensitive film for 7 weeks (D1 receptor), 13 weeks (D2 receptor) or 23 weeks (D3 receptor). Receptor densities were determined by measuring the optical densities of brain regions of interest and comparing them to the standard curve generated by the tritium standards exposed to the same sheet of film (MCID System, Imaging Research Inc., Cambridge, UK). Differences in mean receptor density values between genotypes were analyzed by an unpaired two-tailed Students t-test.

#### Measurements of Serum Amphetamine

For serum amphetamine measurements, blood was collected into serum separation tubes (Starstedt AG & Co.) by retro-orbital bleeds from mice under isofluorane anesthesia, 50 min after their first amphetamine injection. Amphetamine measurements were performed at NMS Labs (Willow Grove, PA) using liquid chromatography followed by mass spectrometry (LC/MS/MS). Serum aliquots of 50 or 100 µl were diluted to 200 µl with human serum. The dilution was taken into consideration to calculate the final concentrations. An internal standard (D5-Amphetamine) and 10% trichloroacetic acid were added to each sample while mixing vigorously. Samples were centrifuged and 200 µl of supernatant from each sample was transferred to autosampler vials for analysis. Samples were analyzed using a Waters Quattro Premier tandem mass spectrometer instrument with electrospray ionization, and a Waters Acquity Ultra Performance LC with an Acquity UPLC HSS T3.1, 2.1 × 50 mm, and 1.8-µm analytical column. Two ion transitions were monitored for amphetamine and the internal standards to assure that there were no interferences. Each analytical run was independently calibrated at concentrations of 5.0, 10, 20, 50, 200, 500 and 1000 ng amphetamine/ml. Controls were run at 30, 375 and 750 ng/ml. During method validation, this LC/MS/MS method had between-run percent CV’s of 6.03, 3.02 and 4.72% at 5.0, 30 and 750 ng/ml, respectively. Amphetamine eluted at approximately 3.5 minutes and the internal standard co-eluted.

#### Open Field Behavior, Rotarod Balance and Fear Conditioning

Horizontal and vertical locomotor activity (distance traveled, rearing) for the evaluation of open-field activity and fear-conditioning tasks was collected via the TruScan System (activity boxes surrounded by 2 rings fitted with infrared sensors, Coulbourn Instrument, Allentown, USA). The same system was used to measure ‘time spent in the margin’ (thigmotaxis) and ‘number of center entries’ for the assessment of anxiety-related behaviors. For fear-conditioning procedures, mice received 5 tones paired with foot shocks (1 per min) in box A after 2 min of free exploration (baseline A) on day 1. On the morning of the following day, mice were placed in the same box without tone or foot shock for 3 min. In the afternoon of the same day, mice were placed in box B (contextually different) and the tone alone was delivered for 3 min after 2 min free exploration of the new box (baseline B). Percent decrease in distance traveled versus baseline in either box was calculated as a measure of association of foot shocks with context or foot shocks with cue. To evaluate motor coordination, mice were placed individually onto an elevated rod accelerating from 4 to 40 rpm over 10 min (Columbus Instruments), and latency to fall was measured.

#### Home Cage Scan

Male wildtype and knockout brothers, 6-7 months of age were single-housed for >7 days prior to videoscanning. Lights were out 7 pm-7 am, and videotaping occurred from 6 pm to 6 am. Data were analyzed by user-trained CleverSys software as described previously (Steele et al., 2007).

#### Social Interaction and Memory Test

According to the method described previously (Crawley, 2000), single-housed male mice were placed individually into a large cage with bedding and two small metal wire enclosures inside, and were allowed to acclimate for 20 min. The male was then removed, and an ovariectomized female was placed into the small wire cage prior to returning the male. The male was videotaped for 2 min prior to removal of the female mouse. After 20 min, the same female was re-introduced for 2 min, and this procedure was repeated four times in total prior to introducing a novel ovariectomized female.

Videotapes were scored, and the amount of time that the male had nose-contact with the cage containing the female was plotted.

#### Olfactory Acuity Test

Wildtype and brother knockout male mice were co-housed and food-deprived by being given access to food for only one hour per day, plus sucrose pellets sprinkled in their home cage (BioServ). Mice were trained for two days by placing individually into a clean cage in the morning and, in the afternoon, were given 15 sucrose pellets on top and beneath the bedding. On the following test day, each mouse went through five trials in which it was removed from the cage, a single pellet was buried, and measurements were taken of the time for the mouse to find the pellet after reintroduction to the cage.

#### Marble Burying

Based on a method described previously (Thomas et al., 2009), male mice were placed into a clean cage and given 30 min to habituate. The mouse was removed, and 9 clean marbles were placed, evenly-spaced, on top of the bedding. The mouse was returned to the cage and videotaped for 15 min. The number of marbles visible at the end was reported as an average across trials. Mice were given three trials across three days.

#### Egocentric and Allocentric T-Maze Tasks

Training was done by an experimenter blinded to genotype, and mice were in mixed-genotype groups. Egocentric T-maze training was conducted in an acrylic cross-shaped maze with white floor and transparent walls. Different departure arms were used so that mice learned to associate the rewarded arm with egocentric cues rather than distal cues. Male mice, food restricted to reach 85% of their free-feeding weight, were given 3-5 habituation sessions in which they were allowed to move freely in the 2 T-maze configurations and to consume chocolate milk placed at the end of the goal arms. Mice were rewarded with chocolate milk for turning in a consistent direction (left vs. right), regardless of their start site and the spatial cues. The rewarded direction was randomly assigned at the beginning of the training unless the mouse had developed a turning bias during the habituation, in which case the opposite direction was baited with food-reward for the training. Each mouse received 10 trials (5 trials from each of 2 start sites), with an inter-trial interval of 30-120 sec, during each daily session for 10 days.

Allocentric T-maze training was conducted in a water maze filled with white dyed water (21°C) into which a mouse was placed at the base of the T to begin the trial. The trial was ended when the mouse touched both forepaws to the submerged escape platform (correct choice) or reached the extremity of the other branch of the T (incorrect choice). The platform was located at a constant position in the experimental room and extra-maze visual cues were provided to instruct the mice of the platform location. Start arms were varied to avert the use of egocentric cues. Percent correct choices and latency to reach one extremity of the T (data not shown) were recorded. Each mouse received 10 trials per day.

#### Step-Wheel Training

All procedures were approved and in accordance with guidelines for the conduct of animal research of Osaka University. The investigator training the mice was blinded to their genotypes. As previously described (Kitsukawa et al., 2011), mice were water-restricted and habituated to run on a step-wheel with moveable pegs in order to reach a water spout. Contact of the paw on the last (12^th^) rung was detected by a voltage change. Mice were trained for 6 days with a regularly-spaced rung-pattern followed by 7 days of training with irregularly-spaced rung pattern 1, followed by 3 days of training with irregularly-spaced rung pattern 2. The whiskers of all mice were then cut off, and the mice were given 2 days in their home cage followed by re-testing for 8 days with irregularly-spaced rung pattern 1. The variance of paw-touch to the 12^th^ rung across training was calculated (Nakamura et al., 2017).

#### Psychomotor Drug Treatments

Different groups of male mice (6-10 months old) were used for each drug treatment. All experiments were conducted genotype-blind. Mice were habituated to injection for 3-5 days and placed in Truscan activity monitors for at least 20 min prior to drug injection. Habituation was used for all drugs with the exception of the haloperidol- and SCH23390-response experiments.

Amphetamine cocaine, SCH23390, SKF83959, SKF81297 and quinpirole were dissolved in saline. SCH23390 and SKF83959 were dissolved in distilled water. Apomorphine was dissolved in 0.1 % acetic acid in water. Haloperidol was dissolved in 1 drop of acetic acid prior to adjusting the volume to the appropriate concentration with saline. All drugs were purchased from Sigma-Aldrich Corp. All drugs were prepared fresh daily, administered at 10 ml/kg body weight and were injected intraperitoneally, except for apomorphine, which was administered sub-cutaneously.

With high or repeated doses of amphetamine, mice show more confined stereotypy and less distance traveled relative to acute or low-dose treatments. We therefore measured both distance traveled and confined stereotypy in CDGI knockout and control mice with repeated administration of low and high doses of amphetamine. For sensitization, mice received daily drug injections for 7 days followed by 7 days of no injection and then a final challenge injection on day 15 as previously described (Crittenden et al., 2014). One exception to this schedule was the 10 mg/kg/day amphetamine experiment in which a two-injection sensitization protocol was used (Valjent et al., 2010).

Distance traveled was computed by infrared photobeam breaks, sampled at 0.5 sec. Stereotypies were measured from 2 min videotapes made at 20 and 50 min post-injection and rated with a keypad scoring system by a rater blinded to genotype as previously described (Crittenden et al., 2014). The highest levels of confined stereotypy occurred around 50 min post-injection for amphetamine. Significance for stereotypy scores was computed by Wilcoxon rank sum test and for distance traveled data by a random effects model. For Figure 2D, the sensitization of the knockouts was normalized to that of wildtypes as follows: the absolute change in distance traveled between day 1 and challenge day for knockouts was divided by the absolute change from day 1 to challenge day in wildtypes, and the ratio was then multiplied by 100 so that the percent change for knockouts could be plotted relative to that for wildtypes. Calculations were based on the peak-response periods videotaped to assess stereotypy, 50-55 min post-injection for amphetamine. From day 1 to challenge day, WT mice showed the expected increase in confined stereotypy and decrease in locomotion, indicating sensitization to the drug. Knockout mice simply maintained their day 1 confined behaviors across all treatment days.

#### Cocaine Intravenous Self-Administration

All procedures were approved by the McLean Hospital Institutional Animal Care and Use Committee. Mice were trained and tested as previously described under an FR 1 schedule of reinforcement in daily 3h sessions, 5-6 days/week (Thomsen et al., 2010). VU0357017 was synthesized at Vanderbilt University (Lebois et al., 2010), dissolved in sterile water (made fresh daily) and administered subcutaneously at 5.6 mg/kg, 15 min before the test session.

#### Corticostriatal Synaptic Plasticity

Experiments were approved and performed in strict accordance with the procedures put forward by the Italian Health Ministry. Corticostriatal EPSPs were evoked by a stimulating electrode placed in cortical regions close to the recording electrode. Bicuculline (10 µM) was added to ∼50% of the experiments to test for contamination of the EPSPs by GABA_A_ receptor-mediated depolarization. The addition of bicuculline did not have an effect on EPSPs, thus data obtained with and without the drug were pooled. For high-frequency stimulation (HFS), 3 stimulus trains were applied (3 sec duration, 100 Hz frequency, 20 sec intervals). HFS protocol was delivered in the presence of 1.2 mM external magnesium to optimize the appearance of LTD. For LTP induction, magnesium was removed from the bathing solution.

#### Single Cell Electrophysiological Recordings

All experimental procedures were performed in accordance with the United States Public Health Service *Guide for Care and Use of Laboratory Animals* and were approved by the Institutional Animal Care and Use Committee at the University of California, Los Angeles (UCLA).

Cells in slices or taken after acute enzymatic dissociation were obtained from 7 wildtype and 10 knockout male mice at 46 and 89 days old (average 68 ± 6 and 68 ± 3 in wildtype and knockout mice, respectively). Whole-cell patch clamp recordings from medium spiny neurons (presumed SPNs) were obtained using standard methods (Cepeda et al., 1998). Cells were identified by somatic size, basic membrane properties (input resistance, membrane capacitance, and time constant), and by addition of biocytin (0.2%) to the internal solution.

Cells in slices or examined after acute enzymatic dissociation were studied using a 40x water immersion lens. The microscope (Olympus BX50WI) was equipped with Nomarski optics and infrared videomicroscopy. The patch pipette (3-5 MΩ) contained the following solution (in mM): Cs-methanesulfonate 130, CsCl 10, NaCl 4, MgCl_2_ 1, MgATP 5, EGTA 5, HEPES 10, GTP 0.5, phosphocreatine 10, leupeptin 0.1 and biocytin (0.2%) (pH 7.25-7.3, osmolality 280-290 mOsm).

Spontaneous postsynaptic currents were recorded in standard ACSF composed of the following (in mM): 130 NaCl, 26 NaHCO_3_, 3 KCl, 2 MgCl_2_, 1.25 NaHPO_4_, 2 CaCl_2_, and 10 glucose, pH 7.4 (osmolality, 300 mOsm). Bicuculline (10 µM) was added to abolish the contribution of spontaneous currents mediated by activation of GABA_A_ receptors. Cells were held at −70 mV to minimize the contribution of GABA_A_ receptors and that of voltage-gated conductances. 4-aminopyridine (4-AP, 100 µM), a K^+^ channel blocker that increases neurotransmitter release, was also applied to the bath after bicuculline blockade to compare EPSC frequencies in cells from wildtype and knockout mice. After characterizing the basic membrane properties of a neuron, spontaneous postsynaptic currents (PSCs) were recorded for variable periods of time (usually 3-6 min). The membrane current was filtered at 1 kHz and digitized at 200 µsec by Clampex (Foster City, CA). Spontaneous PSCs were analyzed off-line using the Mini Analysis Program (Jaejin Software, Leonia, NJ). The threshold amplitude for the detection of an event was adjusted above root mean square noise level (generally ∼5 pA). This software was used to calculate PSCs frequency, amplitude for each synaptic event, and to construct amplitude-frequency histograms. Frequencies were expressed as number of events per second (in Hz). PSCs kinetic analysis used the Mini Analysis Program. Events with peak amplitudes between 10 and 50 pA were grouped, aligned by half-rise time, and normalized by peak amplitude. Events with complex peaks were eliminated. In each cell, all events between 10 and 50 pA were averaged to obtain rise times, decay times, and half-amplitude durations. First- and second-order exponential curves were fitted with a maximum of 5000 iterations, and standard deviations between first- and second-order fits were compared.

Some slices also were used for acute dissociation of neurons. After at least 1 hr of incubation in the oxygenated ACSF, the dorsal striatum was dissected from the coronal slices with the aid of a dissecting microscope. Each striatal slice was placed in an oxygenated cell-stir chamber (Wheaton, Millville, NJ) and enzymatically treated for 20 min with papain (0.5 mg/ml, Calbiochem) at 35°C in a N-[2-hydroxyethyl] piperazine-N-[2-ethanesulfonic acid] (HEPES)-buffered Hank’s balanced salts solution (HBSS, Sigma Chemical) supplemented with (in mM) 1 pyruvic acid, 0.005 glutathione, 0.1 N^G^-nitro-L-arginine, and 1 kynurenic acid (pH 7.4, 300-310 mOsm). After enzymatic digestion, the tissue was rinsed several times with a low Ca^2+^ HEPES-buffered Na-isethionate solution containing (in mM) 140 Na isethionate, 2 KCl, 2MgCl_2_, 0.1 CaCl_2_, 23 glucose, and 15 HEPES (pH 7.4, 300-310 mOsm). Striatal slices were then mechanically dissociated with a series of graded fire-polished Pasteur pipettes. The cell suspension was then plated into a 35-mm nunclon petri dish mounted on the stage of an upright fixed-stage microscope (Zeiss Axioscope, Thornwood, NY) containing a HEPES-buffered salt solution [in mM: 140 NaCl, 23 glucose, 15 HEPES, 2 KCl, 2 MgCl_2_, and 1 mM CaCl_2_ (pH 7.4, 300-310 mOsm)].

Standard whole-cell patch clamp techniques were used to obtain voltage clamp recordings. Electrodes (2.5-3.5 MΩ) were pulled from Corning 8250 glass (A-M Systems, Carlsborg, WA) and heat polished prior to use. The internal solution consisted of (in mM) 175 N-methyl-D-glucamine (NMDG), 40 HEPES, 2 MgCl_2_, 10 ethylene glycol-bis (β-aminoethyl ether)-N, N, N’, N’-tetraacetic acid (EGTA), 12 phosphocreatine, 2 Na_2_ ATP, 0.2 Na_2_ GTP, and 0.1 leupeptin (pH 7.25, 265-270 mOsm). The external solution consisted of (in mM) 135 NaCl, 20 CsCl, 3 BaCl_2_, 2 CaCl_2_, 10 glucose, 10 HEPES, and 0.0003 tetrodotoxin (TTX) (pH 7.4, 300-310 mOsm). Recordings were obtained with an Axon Instruments 200A patch clamp amplifier and controlled by a computer running pClamp (v. 8.01) with a DigiData 1200 series interface (Axon Instruments, Foster City, CA). Data were collected from neurons that had access resistances below 20 MΩ. After whole-cell configuration was reached, series resistance was compensated (70-90%) and periodically monitored. All recordings were made from medium-sized neurons. After obtaining basic membrane properties, NMDA and AMPA (alone or in the presence of cyclothiazide, CTZ 10 µM) were applied for 3 sec duration every 10 sec through a pressure-driven fast perfusion system using an array of application capillaries placed a few hundred micrometers from the cell. To study the modulation of NMDA and AMPA currents, a dopamine D2 receptor agonist, quinpirole (10 µM), was used. A DC drive system controlled by a SF-77B perfusion system (Warner Instruments, Hamden, CT) synchronized by pClamp changed solutions by altering the position of the capillary array. Values for peak currents and peak current densities were calculated for all neurons. Current density was obtained by dividing currents by cell capacitance to normalize values with respect to the size of the cell.

Passive membrane properties of cells in slices or dissociated cells were determined in voltage clamp mode by applying a depolarizing step voltage command (10 mV) and using the membrane test function integrated in the pClamp8 software (Axon Instruments, Foster City, CA). This function reports membrane capacitance (in pF), input resistance (in MΩ or GΩ) and time constant (in µsec or msec). Differences in mean current densities at various voltage commands were assessed with a two-way analysis of variance with one repeated measure followed by multiple comparisons using Bonferroni t-tests. Student t-tests alone were used when only two group means were compared.

#### Fast-Scan Cyclic Voltammetry

All voltammetric experiments were approved and performed in strict accordance with the Institute of Comparative Medicine Laboratory Animal Resources at Columbia University.

Striatal dopamine release was studied in two to five month old male knockout and wildtype mice. Recordings were obtained from the striatum in the first three most rostral coronal slices (300 µm). Three sites in the dorsal striatal region of each slice were measured and averaged. Slices were allowed to recover for 1.5 hr in a holding chamber in oxygenated ACSF at room temperature, and then were placed in a recording chamber and superfused (1 ml/min) with ACSF (in mM: NaCl 125, KCl 2.5, NaHCO_3_ 26, CaCl_2_ 2.4, MgSO_4_ 1.3, KH_2_PO_4_ 0.3, glucose 10) at 36°C.

As described previously (Zhang and Sulzer, 2003), disk carbon fiber electrodes of 5 µm diameter with a freshly cut surface were placed into the dorsal striatum about 50 µm into the slice. For cyclic voltammetry, a triangular voltage wave (−400 to +900 mV at 280 V/s versus Ag/AgCl) was applied to the electrode every 100 ms. Current was recorded with an Axopatch 200B amplifier (Axon Instrument), with a low-pass Bessel Filter setting at 10 kHz, digitized at 25 kHz (ITC-18 board, Instrutech Corporation). Triangular wave generation and data acquisition were controlled by a PC running a house-written IGOR program (WaveMetrics). Striatal slices were electrically stimulated every 2 min with either a single pulse stimulation or a paired stimulus by an Iso-Flex stimulus isolator triggered by a Master-8 pulse generator (A.M.P.I.) using a bipolar stimulating electrode placed ∼100 µm from the recording electrode. Background-subtracted cyclic voltammograms served to identify the released substance. The dopamine oxidation current was converted to concentration based upon a calibration of 5 µM dopamine in ACSF after the experiment.

#### Additive Random Effects Model for Distance Traveled Data

Distance-traveled data were analyzed by a state-space model (Kitagawa, 1996; Smith et al., 2004). Activity monitor data for distance-traveled were chosen for 21 time-points (−15 to 85 min in steps of 5 min) for all wildtype and CDGI knockout mice. Data from five experiments were analyzed, with the numbers of mice in each group ranging from 8 to 12 mice.

We applied this method because we wished to make time-point by time-point comparisons of the group changes in distance-traveled. Independent t-tests comparing groups at each time-point indicated that there was likely to be an effect of genotype, because there were many successive *P*-values that were less than 0.05. However, if corrected for multiple comparisons using Bonferroni, many of these *P*-values were no longer significant. This result was likely due to the fact that the Bonferroni correction does not account for multiple consecutive *P*-values close to zero. Therefore, we turned to a state-space model, which can accommodate point-by-point comparisons and can also be fitted to nonlinear data with peaks at different times. This model assumes that the changes in distance-traveled between time-points are smooth, with the level of smoothing being estimated from the variability of the data. In addition, the model was hierarchical: we estimated the group effects using a Bayesian random effects approach that provided an estimate of the population mean for each group on a given day and with a given drug-dose combination. We then compared the population estimates using Monte Carlo methods. The detailed description of these analyses is as follows.

#### State-Space Random Effects Analysis of Distance Traveled Data

On a given day (day1, day 7 or challenge) and for a given drug-dose combination, the distance-traveled value at time *k* for mouse *j* from the wildtype group is 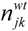. The corresponding data for the knockout mice is 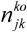.

We assume that all mice from each group have a common time-varying component in their distance-traveled state, *x_k_*, at trial *k*. For the wildtype group, this component follows a random walk as follows:

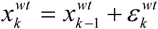 where 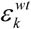 is zero-mean Gaussian with variance 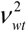. Each mouse *j*’s individual behavioral state is taken to be the same as *x* to within a constant where 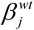 the values 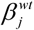 are drawn from a population normal distribution with mean 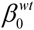 and variance 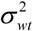. Thus, the distance-traveled individual observations are related to the behavioral state 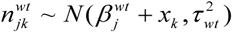 using where 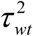 is the group variance estimate and 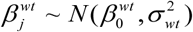. We use an identical model formulation as above for the knockout mice group.

We estimate the population distance-traveled curves for each group by specifying the above Bayesian model in WinBugs (Lunn, 2000). This software estimates the desired posterior distributions using Markov chain Monte Carlo (MCMC). We have used a similar random effects state-space model for animal behavior (Breuer et al., 2005) and the Bayesian estimation method for state-space models (Smith et al., 2009; Smith et al., 2007) in previous reports. In order fully to describe the Bayesian model, we have to specify initial conditions and prior distributions. Specifically, the state is assumed to start at a fixed positive value of 10 (for identifiability, as it is added to a value which is also estimated). Priors on β^wt^ and β^ko^ are uniform on [−1000, 1500]. Priors on precisions are inverse gamma with parameters 0.1 and 0.01 (code given below). Data analyzed with this procedure are shown in Figures 2E and 2F.

We next compared population distributions based on their Monte Carlo samples using the sampling algorithm described in Smith et al. (Smith et al., 2005). Figure S5a-d shows comparison plots of population curves for 8 comparisons across groups and drug treatments. In the bottom left plot of Figure S5a, the black line shows the level of certainty that the wildtype population curve for day 1 is greater than the wildtype population curve for the challenge day 1, i.e., the probability that a sample taken from the day 1 wildtype population is bigger than a sample taken from the challenge day wildtype population. Green asterisks denote locations at which we consider the populations to be very significantly different based even on a Bonferroni corrected *P*-value. Thus, we conclude that the distance-traveled by the wildtype population on day 1 is higher than that on challenge day during 20-35 min post-injection. But the distance-traveled by the wildtype population on day 1 is significantly lower than the distance-traveled on challenge day during the 5 min post-injection time period analyzed.

#### WinBugs Computer Code

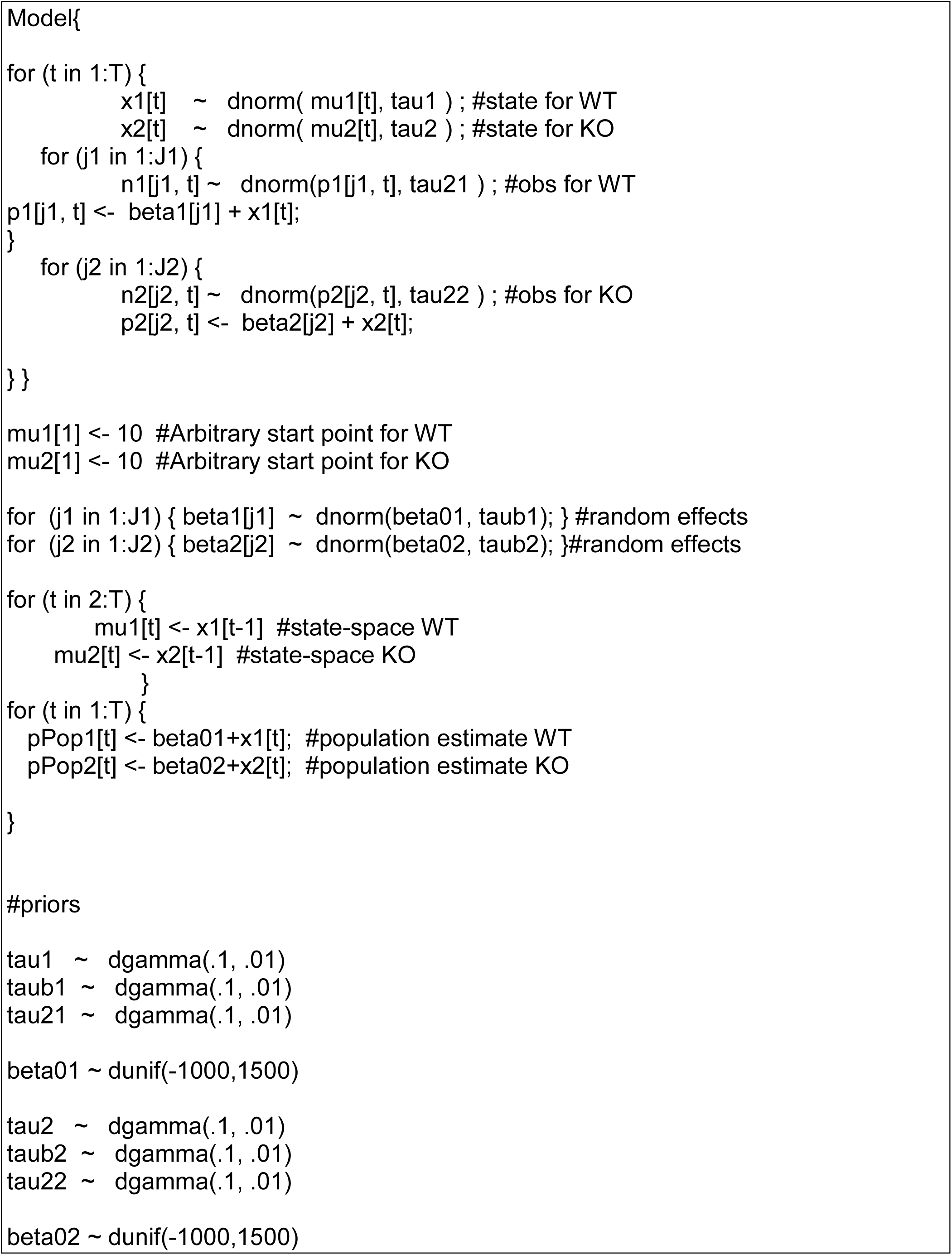

---

**Figure 1S.**
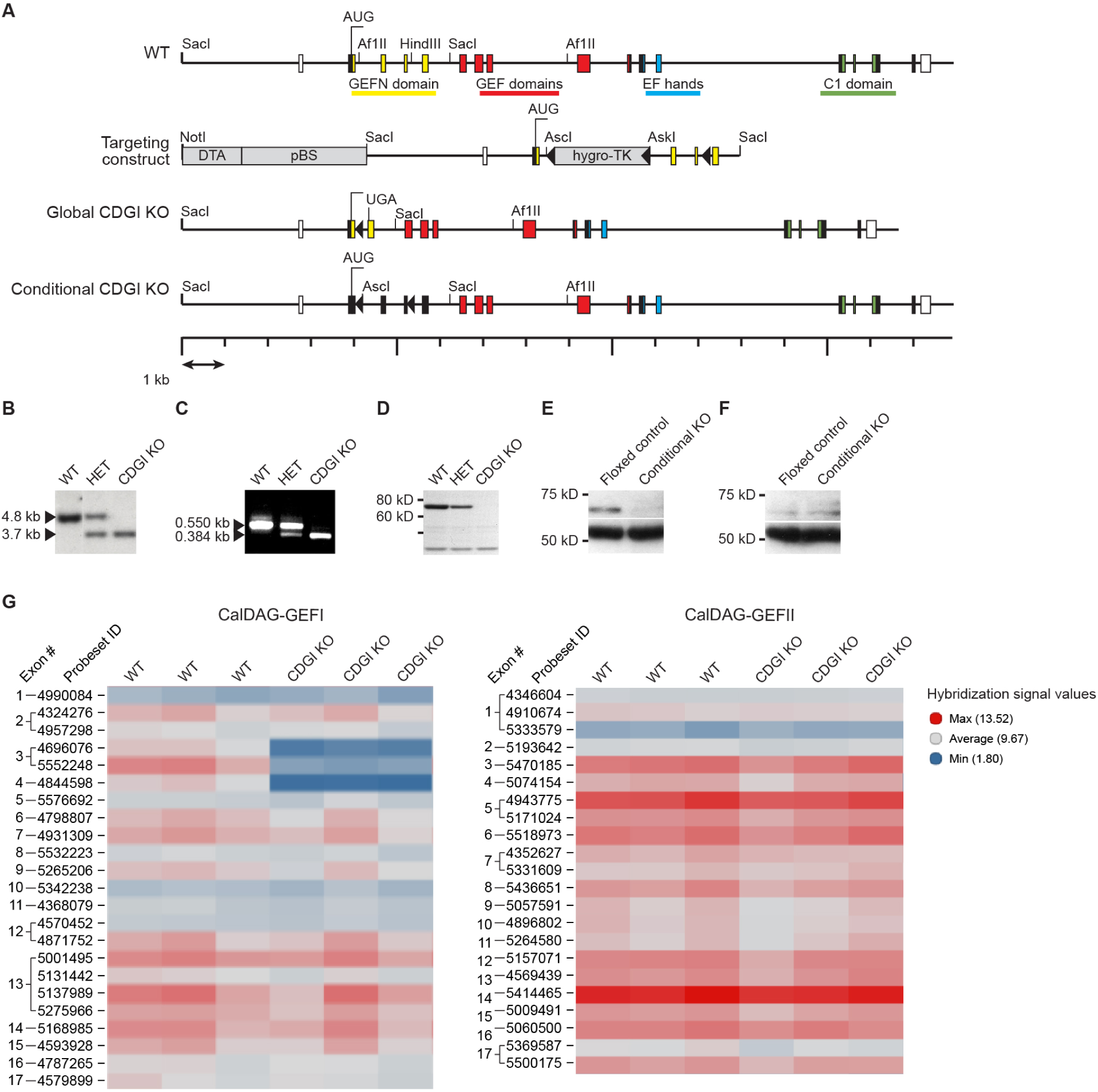
Generation of CDGI Knockout Mice, Related to Figure 1. (A) Diagram of the wildtype (WT) and knockout (KO) gene loci and the targeting construct used to generate KO mice. Colored boxes represent translated exons, and white boxes represent untranslated exons. Colors denote identified sequence domains. Triangles denote LoxP sites. Exons three and four were floxed and deleted by Cre recombinase, either in embryonic stem cells transfected with Cre to create a global knockout, or by crossing to mice with tissue-specific Cre expression to create a conditional knockout. The deletion of exons three and four creates a frameshift mutation and premature stop codon (UGA) in exon five. (B) Southern blots of tail DNA from WT, heterozygous (HET) and global CDGI KO mice confirmed the deletion in CDGI. (C) RT-PCR from global CDGI KO and WT brain RNA demonstrated that the size of the CDGI transcript in mutant mice was reduced as expected. (D) CDGI protein (∼69 kD) was detected in immunoblots of whole brain lysates from WTs and heterozygotes but not in lysates from KOs. (E) Immunoblots of striatal extracts from control mice carrying CDGI^flox/flox^ and conditional KO mice carrying CDGI^flox/flox^ and D1-Cre YAC. (F) Immunoblots show low levels of CDGI expression in neocortical extracts where both CDGI and D1-Cre expression are low. Lower band is a β-tubulin loading control in E and F. (G) CDGI KO mice showed loss of expression from CDGI exons 3 and 4, based on Affymetrix Exon 1.0 microarray data with striatal RNA samples, but no change in the non-deleted exons, and no compensatory changes in expression of the paralog CalDAG-GEFII. *n =* 3WT/3KO.

**Figure S2.**
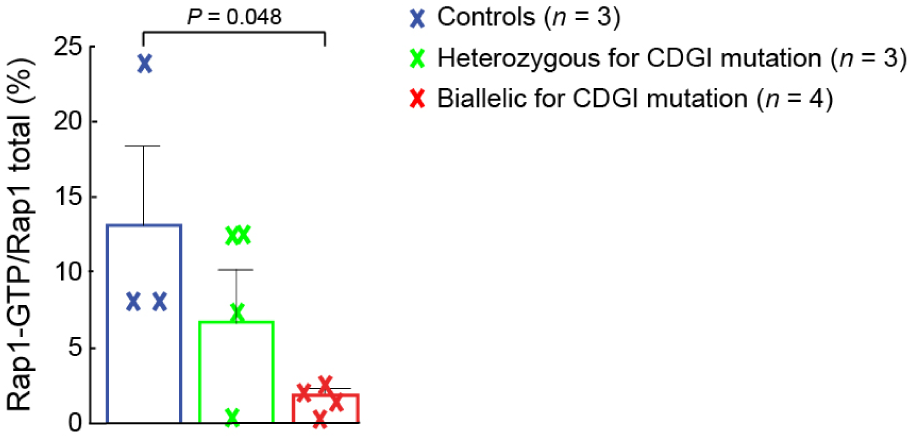
Rap1 Activation Is Reduced in Platelets from Humans with Biallelic CDGI Mutations Relative to Controls, Related to Figure 1. Platelets were activated with ADP (10 µm) for 1 min prior to stopping the reaction and lysing. Lysates were used to prepare immunoblots for Rap1-GTP and total Rap1 and the values plotted on the bar graph reflect the proportion of activated Rap1. *P* value was calculated by two-tailed Student’s t-tests for unpaired samples. See Table S2 for exact values and additional data. Error bars show standard error of the mean.

**Figure S3.**
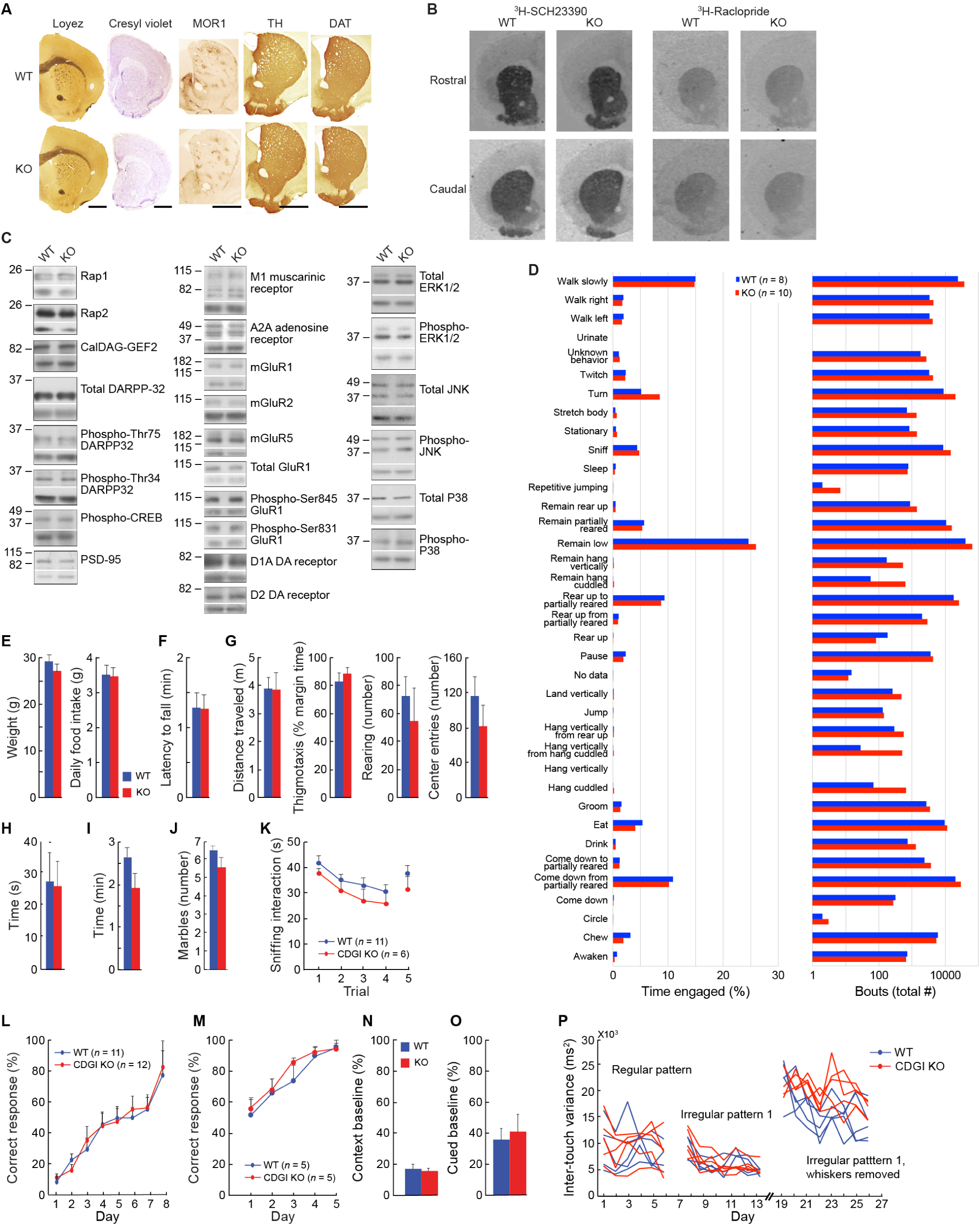
Global CDGI Knockout Mice Have Normal Gross Anatomy of the Brain and Baseline Motor Behaviors, but Abnormal Motor Learning, Related to Figure 2. (A) Gross evaluation of transverse brain hemi-sections did not show differences between global knockout (KO) and wildtype (WT) mice. Sections were stained for white matter (Loyez), cell bodies (Cresyl violet), striosomal immunomarker (Mu-opioid receptor, MOR1), and immunomarkers for dopaminergic terminals (tyrosine hydroxylase, TH and dopamine transporter, DAT). (B) Autoradiograms of rostral and caudal transverse brain sections from global KOs and sibling WTs incubated with tritiated SCH23390 (D1 receptor ligand) and raclopride (D2 receptor ligand) showed equivalent levels of ligand binding between genotypes. See Table S5 for quantitative evaluation. (C) Immunoblots of striatal lysates from WT and global KOs showed grossly equivalent levels of a wide array of neuronal proteins and their phosphoprotein isoforms, as designated to the right of the blot. Location of molecular weight markers is given in kilodaltons to the left of the blot. Bottom bands on each blot show β-tubulin loading controls. Each lane of the blot represents lysates pooled from three mice, and each experiment was performed and evaluated in triplicate. (D) During a 24-hour homecage monitoring period, there were no differences between genotypes in the proportion of total behavioral bouts or number of bouts for any of the automatically detected behaviors (see Steele et al.(Steele et al., 2007) for details of the assay). (E) Adult, age-matched global CDGI KO males and WT brothers had similar weights and total 24-hour plain-chow consumption (*P* = 0.4 for weight and *P* = 0.8 for food consumed by two-tailed Student’s t-tests for unpaired samples; *n=* 12WT, 9KO). (F) KOs and WTs exhibit equivalent latencies to fall from an accelerating rotarod. *P* > 0.05 by two-tailed Student’s t-test for unpaired samples. *n =* 9WT, 11KO. (G) In open field tests, WTs and KOs exhibited equivalent distance traveled (first panel), percent time at field margin (second panel), number of rearing events (third panel), and center entries (fourth panel) during a 60 min period. *P* > 0.05 by two-tailed Student’s t-tests for all measures. *n =* 8WT, 10KO. (H) The latency to retrieve a buried food pellet was not different between WTs and global KOs indicating that olfactory function was not impaired by loss of CDGI. *P* = 0.9 by two-tailed Student’s t-test for unpaired samples. *n =* 7W*T*, 4KO. (I) In a tail-suspension test for learned helplessness (Crawley, 2000), the latency to stop struggling was not different for WT and KO. *P* = 0.1 by two-tailed Student’s t-test for unpaired samples; *n =* 10WT, 7KO. (J) In a 15 min marble-burying test for repetitive behavior(Thomas et al., 2009), the number of marbles remaining unburied, out of 9 total, was equivalent for the two genotypes. *P* = 0.3 by two-tailed Student’s t-tests for unpaired samples; *n =* 9WT, 8KO. (K) In a test for social interaction and memory (Ferguson et al., 2000), WT and global KO male mice sniffed the same female mouse for less time on repeated presentations (trials 1-4) and showed increased interaction when a new female was presented (trial 5). Behavior was not significantly different between genotypes. *P* > 0.19 for genotype comparison on each trial by two-tailed Student’s t-tests. (L) Following the acquisition of the egocentric learning task (shown in Figure 2a), the rewarded arm was switched to test reversal learning, for which WT and CDGI KO mice showed equivalent acquisition. \**P* < 0.05 on all days by unpaired, two-tailed Student’s t-test). (M) CDGI KOs learned to navigate an allocentric, hippocampus-dependent, T-maze at the same rate as sibling controls. Genotype effect: *P* = 0.588, genotype-by-session interaction effect: *P* = 0.713, session effect: *P* < 0.001. (N) In a context-dependent, fear conditioning task, WTs and KOs learned equivalently. Performance is reflected by the reduction in distance traveled in a context paired with foot-shock relative to distance traveled in the context prior to foot-shock. *P* = 0.572; *n* = 7WT, 7KO. (O) WTs and KOs exhibited equivalent fear conditioning in a cued (amygdala-dependent) version of the task as reflected by a similar drop in distance traveled when presented with a tone that was previously paired with foot-shock. *P* = 0.712; *n* = 7WT, 7KO. Means + SEM are shown for E-O. (P) Related to Figure 2b, the performance of individual mice is plotted on the step-wheel task. After whisker-cutting, most CDGI KO mice have a higher variance than control mice for the timing of paw-placement upon the pegs of the wheel.

**Figure S4.**
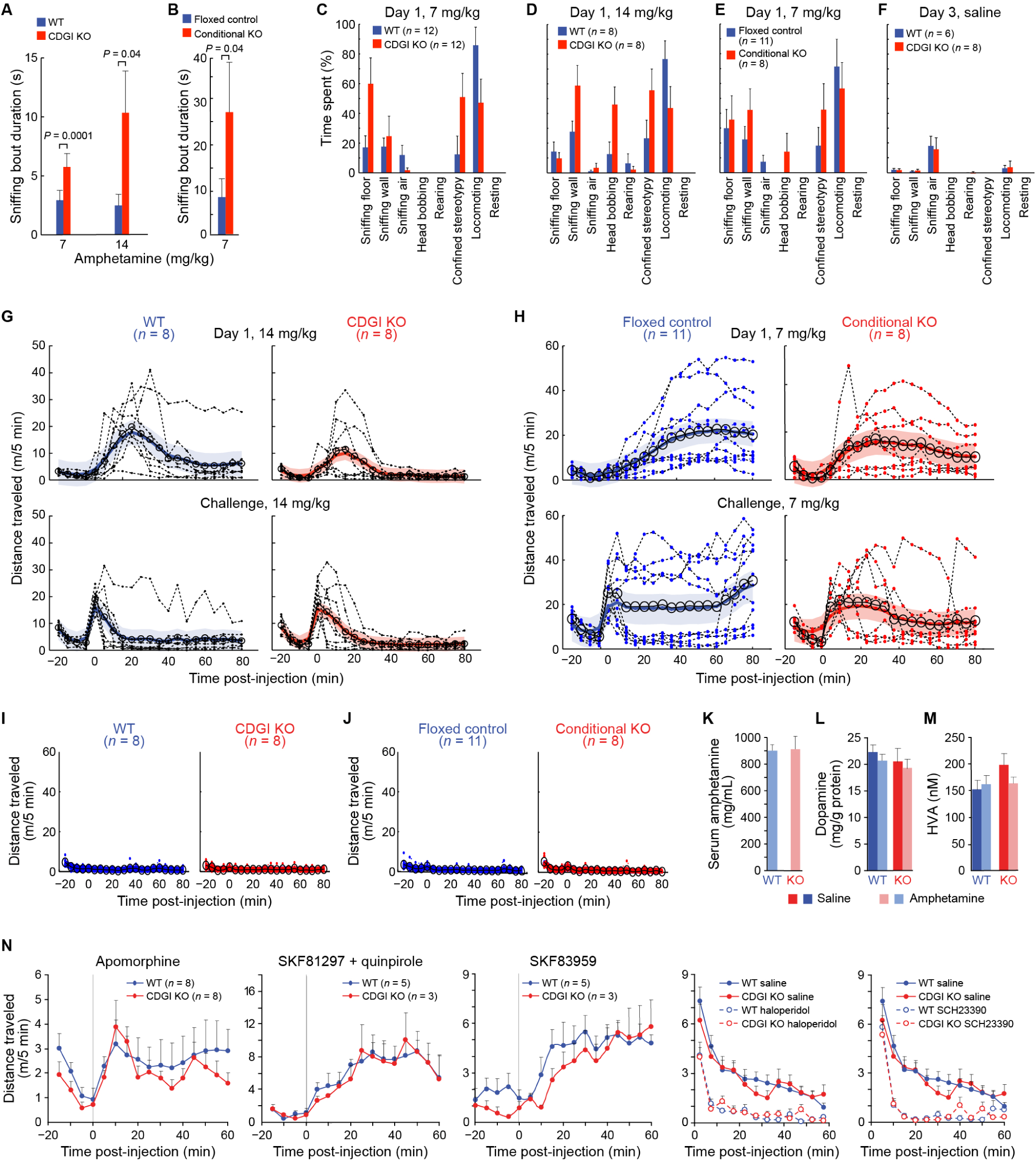
CDGI Knockout Mice Treated with High-Dose Amphetamine Exhibited Significantly Less Distance Traveled than Control Mice in Favor of Increased Stereotypy, but Showed Equivalent Serum Amphetamine, Equivalent Total Striatal Dopamine and HVA, and Equivalent Motoric Responses to Direct Dopamine Receptor Agonists and Antagonists, Related to Figure 2. (A and B) On day 1 of high-dose amphetamine treatment, the average length of each sniffing bout, scored by videotape observation at 50-52 min post-injection, was higher in global CDGI knockout (KO) mice relative to wildtype (WT) sibling controls (A) and in conditional CDGI^flox/flox^ with D1-Cre YAC mice relative to CDGI^flox/flox^ sibling controls (B). *n =* 12KO, 12WT for 7 mg/kg, 8KO, 8WT for 14 mg/kg, and 8 conditional KO, 11 sibling controls for 7 mg/kg. Error bars show standard error of the mean. (C-E) All behaviors that were scored by videotape, at 50-52 min post-injection, showing that both global and conditional KO mice had a tendency for increased stereotypic behaviors (sniffing the wall, confined, head bobbing) relative to the increased locomotor behaviors of their sibling controls. (F) Saline-treated WT and global KO mice did not show differences in scored behavior. (G and H) On day 1 of amphetamine treatment, global KO mice (G) and conditional KO CDGI^flox/flox^ with D1-Cre YAC mice (H) showed less distance traveled than WT or CDGI^flox/flox^ sibling control mice in response to high doses of amphetamine (14 for global KO and 7 mg/kg for CDGI^flox/flox^ mice) on day 1 (top plots), consistent with their increase in stereotypies. WTs showed reduced distance travelled on challenge day (bottom plots) relative to day 1, consistent with stereotypy sensitization. KOs were already in confined stereotypy on day 1 and did not show a significant reduction in distance traveled. Dotted lines are raw data from each mouse; large open circles are population means; colored lines are random effects estimates of the median with 90% confidence intervals. See significance plots in Figure S5. (I and J) There were no significant differences in distance traveled data on saline injection day 3 (injection at time = 0) between global CDGI KO and WT sibling controls (I) or between conditional CDGI^flox/flox^ with D1-Cre YAC mice and CDGI^flox/flox^ sibling (J). See significance plots in Figure S5. (K) Amphetamine levels were equivalent in serum collected from WTs and global CDGI KOs at 50 min post-injection of amphetamine on day 1 (7 mg/kg). *P* > 0.9 by two-tailed Student’s t-test. (*n* = 5WT, 5KO). (L) Total dopamine was measured in striatal homogenates that were collected at 50 min post-injection of amphetamine or saline on challenge day. The challenge injection was given one week after the last repeated saline or amphetamine (14 mg/kg) treatment (see STAR Methods). Dopamine levels were equivalent in WTs and global CDGI KOs, indicating that there was not broad neurotoxicity to dopamine-containing terminals. *n =* 4WT, 5KO for saline and *n =* 8WT, 6KO for amphetamine. (M) COMT activity appeared to be equivalent at 50 min post-injection of saline or amphetamine in WTs and global CDGI KOs, based on measurements of total HVA in tissue collected at 50 min post-injection of amphetamine. *n* = 4 WT + saline, *n* = 7 WT + amphetamine, *n* = 5 KO + saline, *n* = 5 KO + amphetamine. *P* = 0.1 for WT saline vs. amphetamine, *P* = 0.1 for KO saline vs. amphetamine by Student’s unpaired, two-tailed t-tests. (N) Distance traveled for separate groups of mice injected with dopamine receptor agonists and antagonists injected at time 0 in the activity monitors. The combined D1/D2 receptor agonist apomorphine (2.0 mg/kg, s.c.) induced slow locomotion in both WT and KO mice. Mice showed slow locomotor responses to a combined injection of the D1 and D2 agonists SKF81297 (4.0 mg/kg, i.p) and quinpirole (0.025 mg/kg, i.p), and to an injection of the atypical D1 agonist SKF83959 (0.4 mg/kg, i.p.). In mice that were exposed to the activity monitor for the first time, exploratory locomotion was reduced relative to saline-treated controls in both WT and KO mice that were injected with the D2 antagonist haloperidol (1.0 mg/kg, i.p), or the D1 antagonist SCH23390 (0.03 mg/kg, i.p), *n =* 8 global KO, 8WT for each experiment.

**Figure S5.**
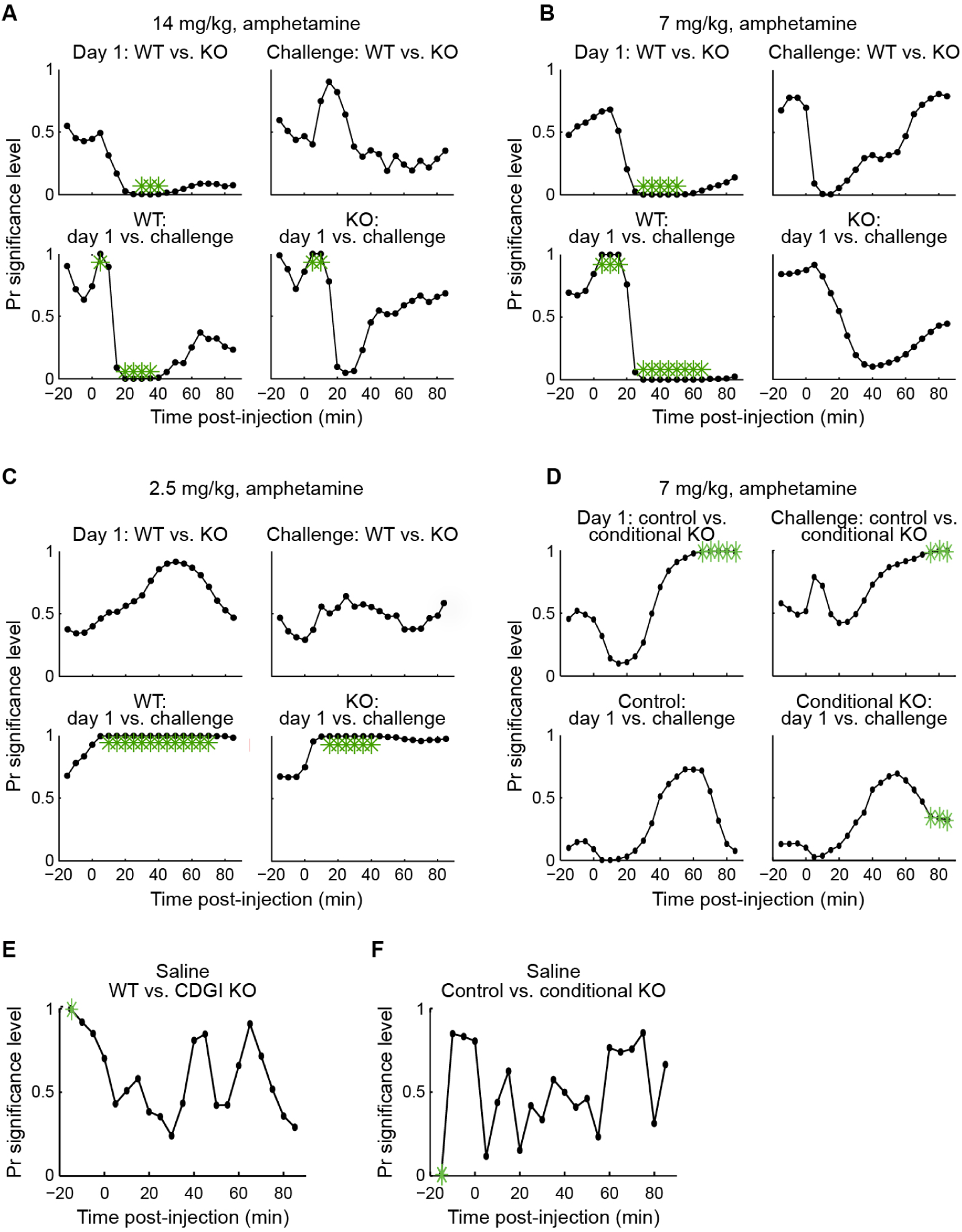
CDGI Knockout Mice Show Significantly Less Distance Traveled after Acute Treatment with High-Dose, but Not Low-Dose, Amphetamine, Related to Figure 2. (A-D) Significance comparisons were made between genotypes on day 1 and challenge day and, to measure sensitization, within genotypes on day 1 vs. challenge day. Plots show comparisons between wildtype (WT) and knockout (KO) mice with 14 mg/kg (A), 7 mg/kg (B) or 2.5 mg/kg (C) amphetamine treatment, and between conditional CDGI^flox/flox^ with D1-Cre YAC mice relative to CDGI^flox/flox^ sibling controls with 7 mg/kg amphetamine treatment (D). Green asterisks indicate significant differences (*P* < 0.05) with Bonferroni corrected comparisons. When the Pr value (Y axes) approaches 0, the first data set (indicated prior to “vs.” in the plot title) is greater than the second data set. For example, the upper left plot of A shows that KOs travelled significantly less than WTs during the 3-50 min post-injection period when stereotypies predominated. Data were analyzed using the state-space random effect model (see STAR Methods). (E and F) Significance plots (equivalent to those shown in A-D) for saline-injected KOs and WTs (E) and for saline-injected conditional KOs and sibling controls (F).

**Figure S6.**
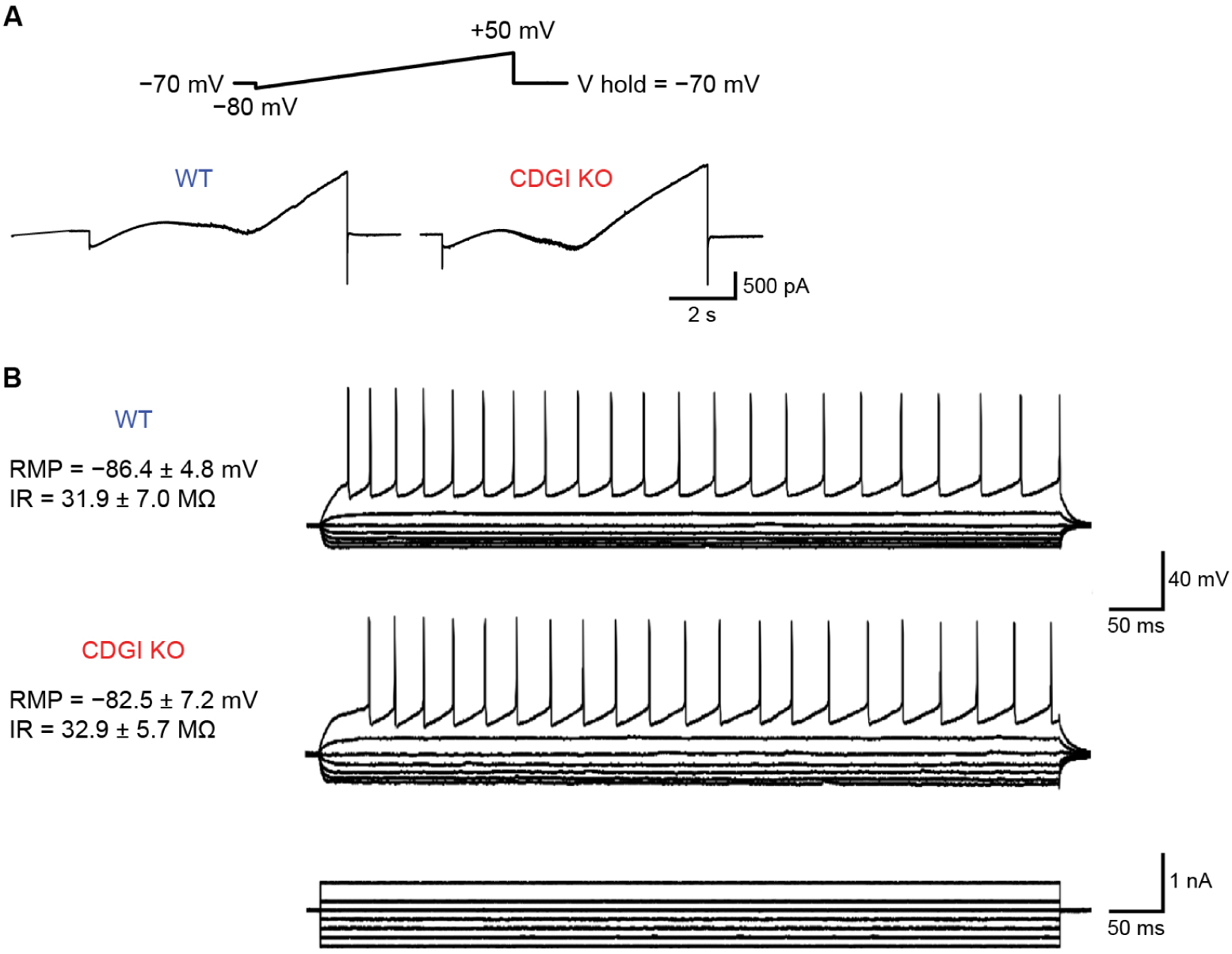
Responses to a Slow Depolarizing Voltage Ramp Command in Cells from KOs and WTs Are Similar, Related to Figure 5E-H. (A) Traces show inward (sodium and calcium) and outward (potassium) currents induced by a depolarizing ramp voltage command from −80 to +50 mV in SPNs from WTs and CDGI KOs. (B) The action potential discharge induced by depolarizing current steps (shown at bottom) was similar in SPNs recorded from WT and KO mice. RMP: resting membrane potential; IR: input resistance, *n =* 8 global KO, 16WT.

**Table S1.** List of All Reported Mutations in CDGI Known So Far, All of Which Are Associated with Defective Hemostasis, Related to Figure 1.

| **Amino acid change**<br>**in protein sequence** | **Variant position in**<br>**coding DNA** | **Variant position in**<br>**DNA relative to NCBI**<br>**sequence**<br>**NM_001098670** | **Reference** |
| --- | --- | --- | --- |
| p.N67Lfs*24 | c.199_200delAA | 386 | 9 |
| p.R113* | c.337C>T | 524 | 2 |
| p.P125* | c.372 -3C>G | 559-3 |  |
| p.F181S | c.542T>C | 729 | 9 |
| p.Q236* | c.887G>A | 893 | 8 |
| p.G248W | c.742G>T | 929 | 4 |
| p.E260* | c.778G>T | 965 | 9 |
| p.Y289C | c.866A>G | 1053 | 9 |
| p.C296R | c.886T>C | 1073 | 9 |
| p.C296Y | c.706C>T | 1074 | 8 |
| p.G305D | c.914G>A | 1101 | 9, 10 |
| p.K309* | c.925A>T | 1112 | 6 |
| p.N330K | c.990C>AorG | 1177 | 9 |
| p.A345P | c.1033G>C | 1220 | 9 |
| p.L360del | c.1078_1080delCTG | 1265 | 6 |
| p.S381F | c.1142C>T | 1329 | 7 |
| p.R494Afs*54 | c.1479_1480insG | 1666 | 9 |
| p.F497Sfs*22 | c.1490delT | 1677 | 9 |

fs: frameshift; \*nonsense mutation that leads to downstream stop codon; del: amino acid deletion.

**Supplementary Table 2.** Platelet Aggregation Assays Show That the Individuals with CDGI Mutations Who Underwent Neurological Testing Show Severely Reduced Platelet Activation as Homozygotes but Not as Heterozygotes, Related to Figure 1.

| Patients<br>Family/<br>subject | Genetics | Sex | Age | Aggregation<br>ADP maximal<br>intensity<br>(10-5 µM) | Aggregation<br>ADP velocity<br>(10-5 µM) | Aggregation<br>TRAP-6 maximal<br>intensity<br>(10-5 µM) | Aggregation<br>TRAP-6 velocity<br>(10-5 µM) | Rap1-GTP/total-Rap1<br>(a.u.) basal | Rap1-GTP/total-Rap1<br>(a.u.) ADP (10 µM) | Rap1-GTP/total-Rap1<br>(a.u.) TRAP-6 (10 µM) |
| --- | --- | --- | --- | --- | --- | --- | --- | --- | --- | --- |
| **Control 1** |  | M | 38 | 86-85 | 161-136 | 85-5 | 77-20 | 0.9 | 7.2 | 46.5 |
| **Control 2** |  | F | 55 | 88-90 | 182-164 | 87-10 | 158-35 | 0.8 | 19 | 4.6 |
| **Control 3** |  | M |  | 88-92 | 171-171 | 88-9 | 104-30 | n.d | n.d | n.d |
| **1.1** | Homozygous<br>G248W | F | 61 | 49-40 | 88-77 | 26-0 | 56-0 | 0.6 | 1.8 | 16 |
| **1.2** | Homozygous<br>G248W | M | 59 | 29-22 | 54-46 | 12-0 | 32-0 | 2.3 | 1.2 | 2.8 |
| **1.3** | Homozygous<br>G248W | M | 55 | 7-0 | 32-0 | 0-0 | 0-0 | n.d | n.d | n.d |
| **2.1** | Heterozygous<br>C296R | M | 47 | 83-60 | 159-123 | 93-8 | 174-30 | 0.8 | 10.6 | 5.4 |
| **2.2** | Heterozygous<br>E260* | F | 46 | 89-85 | 127-112 | 90-2 | 172-2 | 1.2 | 8.8 | 4.2 |
| **2.3** | Double trans-heterozygous<br>C296R E260* | F | 19 | 32-13 | 64-37 | 78-1 | 55-3 | 0.3 | 0.6 | 2.7 |
| **3.1** | Heterozygous<br>N67Lfs*2 | M | 50 | Not possible patient under<br>antiplatelet therapy |  | Not possible patient under<br>antiplatelet therapy |  |  |  |  |
| **3.2** | Heterozygous<br>N67Lfs*2 | F | 54 | 84-82 | 135-119 | 90-1 | 212-10 | 0.6 | 0.1 | 1.4 |
| **3.3** | Homozygous<br>N67Lfs*2 | F | 23 | 16-4 | 38-20 | 79-0 | 81-4 | 0.4 | 0.5 | 1.5 |

**Table S3. Mouse Exon Array, Related to Figure 1**

Provided as an Excel file.

**Table S4. Mouse Total Striatal Amino Acid and Biogenic Amine Measurements, Related to Figure 1**

Provided as an Excel file.

**Table S5.** Dopamine Receptor Ligand Binding, Related to Figure 1.

|  | 3H-SCH23390 (D1) |  |  | 3H-Raclopride (D2) |  |  | 3H-PD128907 (D3) |  |  |
| --- | --- | --- | --- | --- | --- | --- | --- | --- | --- |
|  | WT | KO | *P* | WT | KO | *P* | WT | KO | *P* |
| **DL** | 458 ± 54 | 526 ± 70 | 0.48 | 175 ± 6 | 155 ± 7 | 0.09 |  |  |  |
| **DM** | 455 ± 48 | 505 ± 69 | 0.59 | 138 ± 7 | 124 ± 6 | 0.18 |  |  |  |
| **VL** | 463 ± 48 | 527 ± 62 | 0.45 | 156 ± 8 | 141 ± 8 | 0.25 |  |  |  |
| **VM** | 412 ± 47 | 484 ± 62 | 0.40 | 117 ± 6 | 111 ± 5 | 0.50 |  |  |  |
| **All** | 442 ± 50 | 497 ± 64 | 0.54 | 141 ± 9 | 132 ± 6 | 0.41 | 35 ± 7 | 36 ± 2 | 0.85 |
| **NAc Core** | 352 ± 24 | 370 ± 47 | 0.78 | 65 ± 5 | 58 ± 6 | 0.39 |  |  |  |
| **NAc Shell** | 400 ± 16 | 403 ± 43 | 0.97 | 72 ± 5 | 67 ± 6 | 0.58 |  |  |  |
| **NAc All** | 375 ± 15 | 394 ± 50 | 0.80 | 69 ± 5 | 64 ± 6 | 0.63 | 26 ± 3 | 23 ± 3 | 0.56 |
| **Olf Tubercle** | 397 ± 25 | 384 ± 60 | 0.89 | 85 ± 7 | 75 ± 8 | 0.43 | 31 ± 5 | 29 ± 4 | 0.75 |

Wildtype (WT, *n* = 6) and CDGI knockout (KO, *n* = 8) values were not significantly different by two-tailed Student's t-test for unpaired samples. Means ± SEM are shown. DL: dorsolateral striatum, DM: dorsomedial striatum, VL: ventrolateral striatum, VM: ventromedial striatum, NAc: Nucleus accumbens, Olf: olfactory tubercle.

**Table S6.** Neurological Responses Are Normal in Global CDGI Knockout Mice, Related to Figure 2.

|  | **Wildtype**<br>**(*n* = 7)** | **Knockout**<br>**(*n* = 9)** | **Range** | **Normal**<br>**response** |
| --- | --- | --- | --- | --- |
| **Tremor** | 0 | 0 | 0-1 | 0 |
| **Stereotypy** | 0 | 0 | 0-1 | 0 |
| **Convulsions** | 0 | 0 | 0-1 | 0 |
| **Visual placing** | 3 | 3 | 0-5 | 3 |
| **Grip strength** | 2 | 2 | 0-3 | 2 |
| **Catalepsy** | 2 | 1 | 0-4 | 0 |
| **Wire maneuver** | 0 | 0 | 0-4 | 0 |
| **Flexion reflex** | 2 | 2 | 0-3 | 2 |
| **Turning** | 4 | 4 | 0-4 | 4 |
| **Falling** | 0 | 0 | 0-2 | 0 |

Age-matched males and wildtype sibling were scored according the SHIRPA protocol (Rogers et al., 1997)

**Table S7.** Basic Membrane Properties of SPNs in Slices, Related to Figure 4.

|  | **Cell capacitance**<br>**(pF)** | **Input resistance**<br>**(MΩ)** | **Time constant**<br>**(ms)** |
| --- | --- | --- | --- |
| **Wildtype (*n* = 17)** | 81 ± 3 | 75 ± 8 | 1.5 ± 0.1 |
| **Knockout (*n* = 24)** | 85 ± 4 | 71 ± 4 | 1.5 ± 0.1 |

*P* > 0.05 for inter-genotype comparisons by two-tailed Student's *t*-test for unpaired samples. Means ± standard errors are shown.

**Table S8.**
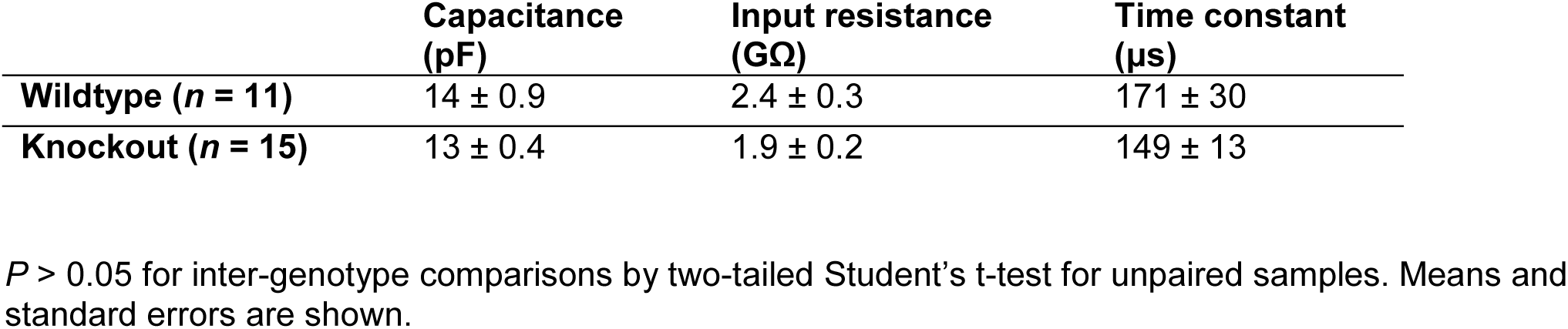
Basic Membrane Properties of Acutely Dissociated Medium Spiny Striatal Neurons, Related to Figure 4.

